# Interplay between β-catenin transcriptional and cell-cell junction activity regulates homeostasis and collective dynamics in the intestinal epithelium

**DOI:** 10.64898/2026.06.10.731321

**Authors:** Thao Nguyen, Sarbari Saha, Fabian Gärtner, Stephan A. Eisler, Jannis Stadager, Dilan Torcuk, Angelika Hausser, Philipp Rathert, Andrew G. Clark

## Abstract

Wnt signaling is crucial for stem cell maintenance during development and homeostasis, and its dysregulation is implicated in cancers across multiple tissues. β-catenin, the primary mediator of canonical Wnt signaling, is also a crucial adaptor protein at cell-cell junctions and is required for maintaining tissue integrity. How the interplay and potential competition between these discrete functions of β-catenin influences tissue homeostasis and organization is not well understood. Here, we address this open question in the intestinal epithelium, which is highly dependent on Wnt signaling during homeostasis and regeneration. Ectopic activation of Wnt signaling in intestinal organoids by pharmacological or genetic perturbation leads to tissue fluidification and loss of epithelial integrity, while disrupting Wnt activity perturbs intracellular β-catenin localization and dynamics at cell-cell junctions. Stabilization of cell-cell junctions increases β-catenin accumulation at junctions at the expense of nuclear localization, while junctional perturbation leads to β-catenin release from junctions into the cytoplasm, leading to dysregulation of Wnt signaling. Together, our findings indicate that in tissues with active Wnt signaling, the interplay between junction- and signaling-associated β-catenin is crucial for stem cell regulation and tissue homeostasis.

## Introduction

Canonical Wnt/β-catenin signaling is crucial for stem cell maintenance and proliferation during development and homeostasis, and changes in Wnt signaling are implicated in cancer in various tissues^1–3^. In addition to its role as a downstream mediator of Wnt signaling, β-catenin plays an essential role as an adaptor protein at adherens junctions (AJs), which mechanically couple the actin cytoskeleton across neighboring cells^4,5^. Modulation of E-cadherin expression or E-cadherin-β-catenin binding elicit phenotypes mimicking Wnt pathway mutants and can lead to intestinal tumor formation, suggesting a competition for these distinct β-catenin functions^6–9^. Cadherins have been proposed to act as a “sink” to prevent β-catenin signaling by preventing nuclear import, though this effect has been suggested to be independent of cell-cell adhesion^10–12^. In other contexts, the signaling and adhesion roles of β-catenin appear to be independent^13–16^. These conflicting reports suggest that the potential interactions of these distinct β-catenin functions may be cell- and tissue-type dependent, which could arise from the specific cellular requirement of Wnt signaling. Indeed, previous reports have suggested that cells with active Wnt signaling are sensitive to β-catenin sequestration by E-cadherin, while E-cadherin is not a major signaling regulator in cells with inactive Wnt signaling^12^. Stem cell proliferation and differentiation in the mammalian intestinal epithelium is highly dependent on Wnt signaling during homeostasis and regeneration. However, it is unclear how this activity is regulated by the interplay between the signaling and adhesion functions of β-catenin.

The small intestine is lined by a simple polarized epithelial layer that is compartmentalized into crypts and villi. Crypts contain stem and progenitor cells that rapidly divide in the transit amplifying (TA) zone before fully differentiating and actively migrating toward the villus tip, with the exception of Paneth cells. The migratory differentiated cells are extruded into the lumen 3-5 days after exiting the crypt^17–19^. Intestinal stem cells maintain their proliferative and self-renewing capacity through morphogen signaling, most prominently via canonical Wnt/β-catenin signaling. A sharp gradient in Wnt signaling at the crypt base is regulated by secretion of Wnt ligands by Paneth cells as well as by mesenchymal cells in the underlying *lamina propria*^20–22^. In the absence of Wnt ligands, intestinal stem cells spontaneously differentiate, resulting in a loss of intestinal stem cells and a collapse of homeostasis^20^. In the presence of Wnt ligand, canonical Wnt signaling is regulated by the β-catenin “destruction” complex, which contains Adenomatous Polyposis Coli (APC), Glycogen Synthase Kinase 3β (GSK-3β) and Casein Kinase 1 (CK1) as well as the scaffolding protein Axin^23^. Phosphorylation of β-catenin by GSK-3β and CK1 targets β-catenin for ubiquitination, leading to its proteolytic degradation^9^. During canonical Wnt signaling, the binding of Wnt ligand to Frizzled (Fzd) receptors on stem cells leads to the repression of the destruction complex, resulting in the accumulation of β-catenin, which is then transported to the nucleus. Once in the nucleus, β-catenin binds and activates T-cell factor/Lymphoid enhancer factor (TCF/Lef), which promotes transcription of downstream genes associated with increased proliferation, stemness and survival^24^. How the intracellular localization dynamics of β-catenin are regulated, and how this impacts downstream signaling, is currently not well understood.

Colorectal cancer (CRC) is primarily initiated by mutations in the Wnt signaling pathway, most commonly APC loss-of-function mutations in Lgr5+ stem cells at the crypt base^25^. Familial adenomatous polyposis (FAP), a hereditary condition most often associated with mutations in APC, results in a near 100% incidence of CRC, typically with early onset^26^. APC mutations lead to epithelial dysplasia; subsequent activation of oncogenes (*KRAS*, *BRAF)* and loss of tumor suppressors (*P53, DCC, PTEN*) leads to invasive carcinoma^27–29^. Re-introduction of functional APC can reverse CRC progression, even in invasive carcinomas harboring *KRAS* and *P53* mutations^30^. APC knock-out in mice leads to mis-localization of Paneth cells from the crypt base to being scattered along the crypt-villus axis^31^. This loss of normal cell patterning suggests that the β-catenin activity is important not only for maintaining the stem cell niche, but also impacts the overall organization of the intestinal epithelium. Modulation of cell-cell junction complexes results in changes in collective cell dynamics and tissue organization^32–34^. This suggests that tumor initiating mutations in APC could drive tissue dysplasia by directly modulating cell-cell junctions, though this has not been directly addressed.

In this study, we investigate the interplay between the Wnt signaling and cell-cell junction functions of β-catenin in the small intestine. Combining quantitative immunostaining, biochemical analysis and photoactivatable live imaging probes in intestinal organoids, we find that perturbing Wnt signaling regulates intracellular β-catenin localization patterns and dynamics at cell-cell junctions. On the tissue scale, increasing Wnt signaling induces tissue fluidification, while reducing Wnt signaling leads to spontaneous stem and progenitor cell differentiation and jamming-like behavior, concomitant with changes in cell morphology. Together, these changes lead to a loss of epithelial barrier function, suggestive of dysregulated cell-cell adhesion integrity. Stabilization or abrogation of cell-cell junctions disrupts normal tissue patterning and Wnt signaling dynamics, further indicating cross-talk between the signaling and adhesion roles of β-catenin. This phenocopies the disruption of Wnt signaling following pharmacological and genetic perturbations, which leads to a dramatic loss of homeostatic tissue organization, including changes in collective cell dynamics and tissue-scale cellular compartmentalization. Taken together, our evidence suggests that in the intestinal epithelium, a competition between the signaling and cell-cell junction functions of β-catenin is critical for regulating downstream Wnt signaling in stem cells and homeostasis.

## Results

### Wnt signaling regulates the intracellular localization of β-catenin

Disruptions in Wnt signaling lead to abnormal tissue organization and tumorigenesis in the intestine^25,31^. This loss of physiological tissue organization could be mediated changes in cell-cell adhesion due to dysregulation of β-catenin localization and activity at cell-cell junctions. To first establish a quantitative baseline of intracellular β-catenin localization patterns, we immunostained for β-catenin in mouse small intestinal tissue (Figure 1A, top row). We next segmented individual cells in 2D confocal slices from histological samples and analyzed the relative localization of β-catenin at cell-cell junctions. To map junctional β-catenin levels with respect to cell position in the tissue, we using a normalized positional scale from the crypt bottom to the villus, where × = 0 corresponds to the crypt base, × = 1 corresponds to the top of the crypt and × > 1 indicates positions along the villus. The relative intensity of junctional β-catenin was lowest at the crypt base and gradually increased before reaching a plateau along the villus (Figure 1B, top row). In contrast, E-cadherin accumulation at cell-cell junctions was constant along the crypt-villus axis.

**Figure 1.**
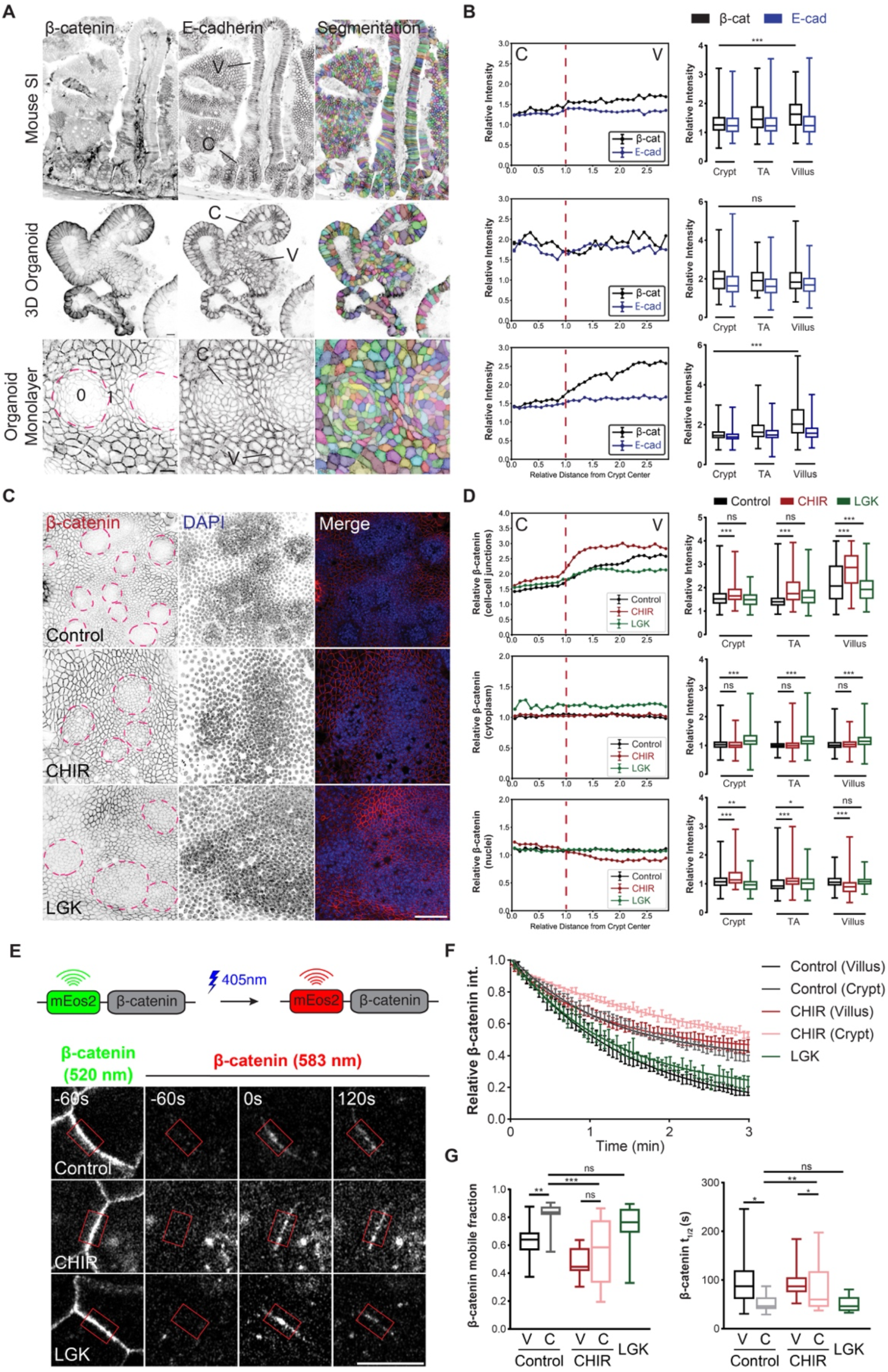
Wnt signaling regulates the intracellular localization and dynamics of β-catenin. **A.** Representative micrographs of mouse small intestinal tissue (top row), 3D organoids (middle row) and organoid monolayers (bottom row) immunostained for β-catenin and E-cadherin and segmented. Magenta dashed lines indicate the crypt-like compartments in organoid monolayers, and lines indicate crypt-like (C) and villus-like (V) compartments. SB: 50µm. **B.** *Left*, Binned linescans of junctional β-catenin and E-cadherin intensity (relative to cytoplasm intensity) of single cells along a computationally reconstructed crypt-villus axis for conditions in *A*. Position values along the crypt-villus axis are rescaled such that the crypt center is at position × = 0, the crypt edge at position × = 1, and the villus-like compartment at positional values × > 1. Binned intensity values indicate mean ± SEM. *Right*, boxplots of relative intensity of β-catenin and E-cadherin of individual cells from crypt, transit amplifying (TA) and villus compartments. Data compiled from *n* =9, 9, 9 organoid monolayer and *N* = 3, 3, 3 independent experiments. **C.** Representative micrographs of organoid monolayers treated with CHIR or LGK and immunostained for β-catenin and counterstained using DAPI. Magenta dashed lines indicate the crypt-like compartments. SB: 50µm. **D.** *Left*, Binned linescans of relative β-catenin intensity in different intracellular compartments (cell-cell junctions, cytoplasm, nuclei) along the reconstructed crypt-villus axis for conditions in *C*. *Right*, boxplots of relative intensity of β-catenin in different intracellular compartments of individual cells from crypt, transit amplifying (TA) and villus compartments. The relative intensity of each cellular compartment is normalized with cytoplasmic signal. Data compiled from *n* = 9, 9, 9 and *N* = 3, 3, 3 independent experiments. **E.** Representative timelapse montage of fluorescence loss in photoactivation (FLIP) experiment using the photo-switchable probe mEos2-β-catenin signal. Cells in organoid monolayers treated with CHIR or LGK were photoactivated at t = 0 in the indicated region of interest (ROI) at cell-cell junctions (red boxes). **F.** Quantification of mean fluorescence of mEos2-β-catenin (583 nm) in the ROI after photoactivation within crypt-like (C) and villus-like (V) domains over time or following CHIR or LGK treatment. **G.** Boxplots of mobile fraction and decay half-time (t_1/2_) extracted from traces used to compile mean data in *F*. Data compiled from one cell-cell contacts from *n* = 30, 30, 28 cell-cell contacts and *N* = 3, 3, 3 independent experiments (same data used for line plot in *F*). Statistical tests performed using one-way ANOVA tests (p<0.001 for all plots) and Bonferroni’s post-hoc test (*p<0.05, **p<0.01, ***p<0.001, ns:p>0.05).

We next investigated whether a similar gradient in junction-associated β-catenin was also present in 3D organoids by immunostaining for β-catenin and E-cadherin (Figure 1A, middle row). To obtain better intracellular resolution while maintaining distinct crypt-villus structures, we generated organoid monolayers, which self-organize into distinct crypt-like and villus-like compartments and display homeostatic cellular turnover^35^. We then immunostained organoid monolayers for β-catenin and E-cadherin and segmented individual cells and intracellular compartments in 3D (Figure 1A, bottom row). To analyze intracellular localization along the crypt-villus axis in organoid monolayers, we assigned each segmented cell a normalized position relative to the center of the nearest crypt and binned the data according the relative position, where × = 0 corresponds to the crypt-like compartment center, × = 1 corresponds to the outer contour of the crypt-like compartment and × > 1 indicates positions in the villus-like regions outside of the crypt-like compartment. This allowed for a reconstruction of the crypt-villus axis from the randomly positioned crypt-like compartments in organoid monolayers to compare with the small intestine and 3D organoid immunostainings. While 3D organoids displayed a noisy but constant junctional accumulation of β-catenin along the crypt-villus axis, organoid monolayers exhibited a β-catenin gradient similar to the native intestinal tissue, with lower β-catenin in the center of the crypt-like compartment and higher intensities in the villus-like compartment (Figure 1B, middle/bottom rows). These data suggest that β-catenin localization varies along the crypt-villus axis and that organoid monolayers better captures this feature of the native tissue compared to organoids cultured in 3D matrices.

We next sought to determine how perturbation of Wnt signaling activity influences the intracellular β-catenin localization gradient along the crypt-villus axis. To this end, we pharmacologically enhanced Wnt signaling in mouse small intestinal organoids by treating with CHIR99021 (CHIR), a GSK-3 inhibitor, or repressed Wnt signaling by treating with LGK974 (LGK), an inhibitor of Porcupine (PORCN), which is required for Wnt ligand release from Paneth cells^36^. We first validated the effects of CHIR and LGK on transcription of Wnt pathway components in 3D organoids by RT-qPCR (Figure S1A). Next, we treated organoid monolayers with CHIR or LGK, immunostained for β-catenin and segmented nuclei and cells in 3D from confocal imaging stacks (Figure 1C). CHIR treatment resulted in increased junctional β-catenin in both the crypt- and villus-like compartments as well as the transit amplifying (TA) compartment, while LGK treatment caused a decrease in junctional β-catenin specifically in the villus-like compartment (Figure 1D). In contrast, CHIR treatment had no effect on cytoplasmic levels of β-catenin, while LGK treatment increased cytoplasmic β-catenin in all compartments. Nuclear β-catenin increased in the crypt-like and TA compartments following CHIR treatment, but decreased slightly in the villus compartment. LGK treatment induced a slight reduction in nuclear β-catenin in the crypt-like compartment.

Thes observations suggest that there may be competition for β-catenin localization at cell-cell junctions and the nucleus, and that intracellular localization can be regulated by upstream Wnt pathway activity. To further validate these changes in intracellular β-catenin localization following perturbation of Wnt signaling, we performed cellular fractionation and Western blot analysis (Figure S1B). Cytoplasmic and nuclear fractions were validated by the presence of GAPDH and Lamin A/C, respectively. While nuclear accumulation of β-catenin was largely unaffected by CHIR or LGK treatment, cytoplasmic levels of β-catenin were highly elevated following CHIR treatment, regardless of posttranslational modification (Figure S1C). These data indicate that β-catenin nuclear import is an active process that can be regulated independently of cytoplasmic β-catenin accumulation.

Recent work has suggested the dynamic nucleoporin protein NUP98 is involved in β-catenin nuclear import during Wnt signaling in colorectal cancer^37^. To determine how Wnt signaling influences NUP98 localization, we treated wt organoids in 3D and monolayer culture formats with CHIR or LGK and immunostained for NUP98 (Figure S1D). Quantification of 3D segmentations of individual cells and nuclei in organoid monolayers revealed that NUP98 accumulation at the nuclear cortex increased following CHIR treatment but was unaffected by LGK treatment (Figure S1E). To determine how NUP98 protein levels are regulated by Wnt signaling, we treated organoids with CHIR or LGK and extracted total RNA and protein lysate. Western blot analysis revealed that overall levels of both β-catenin and NUP98 increased following CHIR treatment and decreased following LGK treatment (Figure S1F,G). Taken together, our results suggest that Wnt signaling in intestinal epithelial cells influences β-catenin localization at cell-cell junctions and can also regulate expression and localization of nucleoporins involved in β-catenin nuclear import.

### Wnt signaling regulates junctional β-catenin dynamics

The intracellular localization of β-catenin is regulated by its binding dynamics in different cellular compartments. The differences in junctional localization following CHIR and LGK treatment may thus reflect changes in β-catenin dynamics at cell-cell junctions. To investigate this directly, we virally transduced organoid monolayers with murine β-catenin fused to the photoactivatable fluorophore mEos2 and performed fluorescence loss in photoactivation (FLIP)^38^. Prior to photoactivation, mEos2-β-catenin accumulated at junctions and was visible only in the green spectrum (emission peak: 520 nm; Figure 1E; Video S1). Following activation in regions of interests (ROIs) at cell-cell junctions, mEos2-β-catenin became visible in the red spectrum (emission peak: 583 nm). Quantifying mean fluorescence in the red spectrum in each ROI over time, we found that the activated pool of mEos2-β-catenin decreased according to an exponential-like decay (Figure 1F). We next fit these data to exponential functions to extract the mobile fraction and decay half-time (t_1/2_; Figure 1G). In untreated control organoid monolayers, β-catenin turnover at junctions in cells was significantly faster in the crypt-like domain, with a mobile fraction of ∼85% (compared with ∼65% for cells in the villus-like compartment) and a t_1/2_ of ∼45 sec (compared with ∼90 sec for cells in the crypt-like compartment). Wnt activation via CHIR treatment led to a slower β-catenin dynamics in both crypt- and villus-like compartments, with a decreased mobile fraction and increased t_1/2_. LGK treatment, conversely, led to increased mobility of β-catenin, and crypt-like domains were no longer present in LGK-treated monolayers. Junctional β-catenin dynamics in these monolayers was most similar to the villus-like domains of untreated crypts, with a relatively high mobile fraction and low t_1/2_. Together, these data suggest that increased Wnt signaling in intestinal epithelial cells reduces β-catenin recycling from cell-cell junctions, implying cross-talk between the signaling and junctional roles of β-catenin and correlating faster junctional turnover dynamics with increased junctional accumulation.

### Modulation of Wnt signaling perturbs intestinal organoid dynamics and barrier function

Our results indicate that perturbation of upstream Wnt signaling leads to changes in the localization and dynamics of junctional β-catenin, an essential component of cell-adhesions. Such disruptions in cell-cell junction complex formation are likely to influence tissue-scale collective migration patterns. To test this, we cultured monolayers using organoids expressing membrane-targeted tdTomato (mT)^39^. Following treatment of organoid monolayers with CHIR or LGK, we performed live confocal microscopy and analyzed the dynamics of cells in crypt-like and villus-like compartments using particle image velocimetry (PIV; Figure 2A; Video S2). CHIR-treated organoid monolayers exhibited a ∼3.5-fold increase in migration velocity within the crypt domain, in addition to a slight increase in correlation length (Figure 2B). LGK treatment led to a minor decrease in correlation length, with no significant effect on migration velocity in crypt-like domains. Analyzing migration dynamics in the villus-like compartment, we found that both CHIR and LGK treatment led to a minor, but significant increase in mean velocity. CHIR-treated monolayers additionally displayed a minor increase in correlation length, while LGK-treated monolayers migrated with a ∼4-fold increase in correlation length compared with untreated controls.

**Figure 2.**
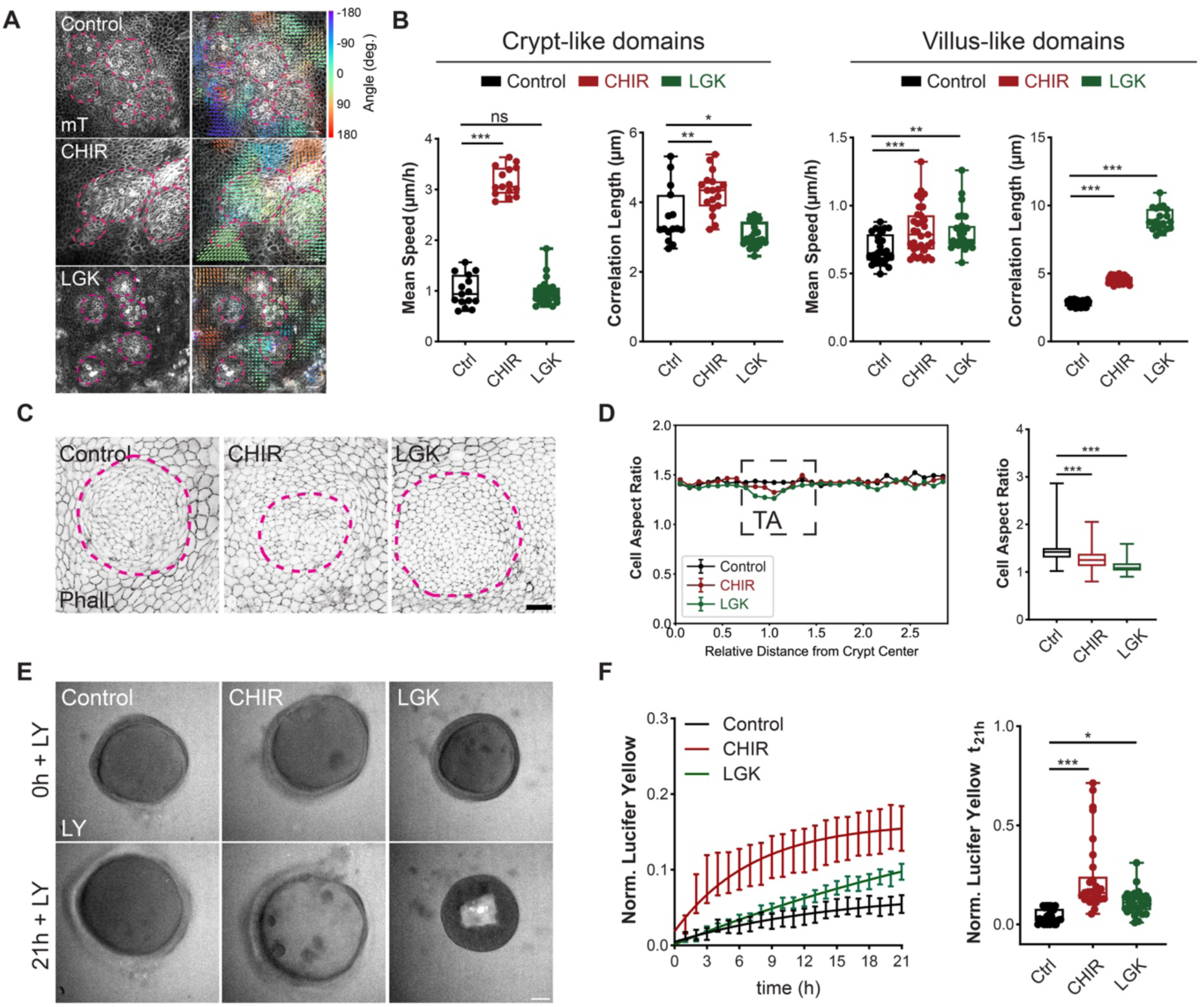
Wnt signaling regulates intestinal tissue dynamics and cell morphology to maintain barrier function. **A.** Representative brightfield micrographs and particle image velocity (PIV) analysis of monolayers generated from organoids stably expressing membrane-targeted tdTomato (mT) treated with CHIR or LGK. SB:50 µm. Scale vectors: 12 μm/hr. **B.** Quantification of mean instantaneous speeds and correlation lengths for crypt-like and villus-like domains of organoid monolayers analyzed by PIV as in *A.* Each dot represents one organoid monolayer from *n* = 16, 18, 18 and *N* = 3, 3, 3 independent experiments. **C.** Representative micrographs of organoid monolayers treated with CHIR or LGK and stained with Phalloidin (Phall.) for F-Actin. Magenta dashed ring indicates crypt-like domains. SB: 50 μm. **D**. Quantification of cellular aspect ratio of individually segmented cell within monolayer as a function of position relative to the crypt center. *Left*, binned averages (mean ± SEM) of cellular aspect ratio at relative distances from the crypt center. Box with black dotted line indicates the transit amplifying (TA) zone. Relative distances to the crypt are rescaled such that the crypt center is at position 0, the crypt edge at position 1, and the villus-like compartment at values >1. *Right*, boxplot of cellular aspect ratios in the TA zone for conditions as in *C*. Data compiled from *n* = 10, 9, 9 organoid monolayers and *N* = 3, 3, 3 independent experiments. **E.** Representative micrographs of 3D intestinal organoids treated with CHIR or LGK, after adding Lucifer Yellow (LY) to the organoid medium. SB: 50µm. **F.** *Left*, Quantification of relative luminal fluorescence (organoid lumen to medium fluorescence intensity) over time after adding LY. Bar and whiskers indicate mean ± SEM, solid line indicates exponential fit. *Right*, quantification of normalized luminal LY intensity after 21 h for condition as in *E*. Each dot represents one organoid from *n* = 40, 45, 42 and *N* = 3, 3, 3 independent experiments. Statistical tests performed using one-way ANOVA tests (p<0.001 for all plots) and Bonferroni’s post-hoc test (*p<0.05, **p<0.01, ***p<0.001, ns:p>0.05).

High correlation lengths are indicative of solid-like behavior, or tissue “jamming”^40,41^. Tissue jamming is associated with more well-packed tissues with isotropic hexagonal cell shapes^42^. To investigate whether increased correlation length also correlated with reduced cellular elongation, we segmented individual cells in CHIR- and LGK-treated monolayers and measured the in-plane aspect ratio (AR; Figure 2C). Reconstructing the crypt-villus axis from cell segmentations, we found that cells localized specifically in the transit amplifying (TA) zone displayed reduced AR following CHIR and LGK treatment (Figure 2D). Taken together, these data suggest that CHIR treatment, and to a much larger extent, LGK treatment, promotes a more solid-like epithelial monolayer, which could reflect changes not only in Wnt signaling, but also in cell-cell adhesions due to modifications in β-catenin localizations or activity.

Stable cell-cell junctions are required for maintaining epithelial barrier integrity, a crucial function of the intestinal epithelium to prevent infection of potential pathogens. As perturbation of Wnt signaling disrupts cell migration patterns, this could also therefore influence barrier integrity. To investigate this, we employed a live-imaging-based organoid permeability assay where the small fluorophore Lucifer Yellow (LY) is added to the organoid medium^43^ (Figure 2E; Video S3). To determine epithelial permeability, we quantified relative luminal LY fluorescence over time (Figure 2F). Consistent with the increase in cell migration and tissue fluidity during CHIR-treatment, we observed an influx of LY into the organoid lumen following CHIR-treatment that was significantly higher compared to untreated control organoids. We also observed a moderate increase in luminal fluorescence in LGK-treated organoids, which likely reflects autofluorescence in the lumen due to excess cell extrusion into the organoid lumen^43^. Together, these experiments suggest that perturbation of Wnt signaling compromises intestinal epithelial dynamics, organization and barrier function.

### Perturbation of cell-cell junctions dysregulates Wnt signaling and epithelial homeostasis

Consistent with β-catenin’s role as an adaptor protein at cell-cell junctions, we observe rapid changes in collective migration dynamics and barrier function following acute pharmacological Wnt perturbation. To further investigate the interplay between signaling and junctional functions of β-catenin during intestinal homeostasis, we next probed how the modulation of cell-cell junctions influences tissue organization, dynamics and signaling. To this this end, we stabilized cell-cell junctions using an intestinal organoid line with a mild overexpression of E-Cadherin^44^ (mTmG/Cdh1-GFP, henceforth referred to as mTmG/E-cad-GFP; Figure S2A,B). To destabilize cell-cell junctions, we treated mTmG organoid monolayers with a functional antibody targeting the extracellular domain of E-cadherin (ECCD-1), which disrupts cadherin binding^45^. We next generated intestinal organoid monolayers from mTmG/E-cad-GFP organoids or parental mTmG organiods with or without ECCD-1 treatment and immunostained for β-catenin and APC (Figure 3A). We then quantified β-catenin and APC localization along the spatially reconstructed crypt-villus axis. In cells in the crypt-like domain, the relative accumulation of β-catenin at cell-cell junctions was higher in mTmG/E-cad-GFP monolayers, while nuclear accumulation was lower (Figure 3B). ECCD-1 treatment showed the opposite effect, with reduced junction-associated β-catenin and increased nuclear β-catenin. These results suggest that stabilization of cell-cell junctions leads to increase β-catenin recruitment at junctions at the expense of nuclear β-catenin, while destabilization of junctions results in release of β-catenin from junctions and a corresponding increase in nuclear β-catenin. APC expression increased along the reconstructed crypt-villus axis and was also influenced by junctional stability; E-cadherin overexpression resulted in reduced APC expression across all compartments, while ECCD-1 treatment led to increased APC expression (Figure 3C).

**Figure 3.**
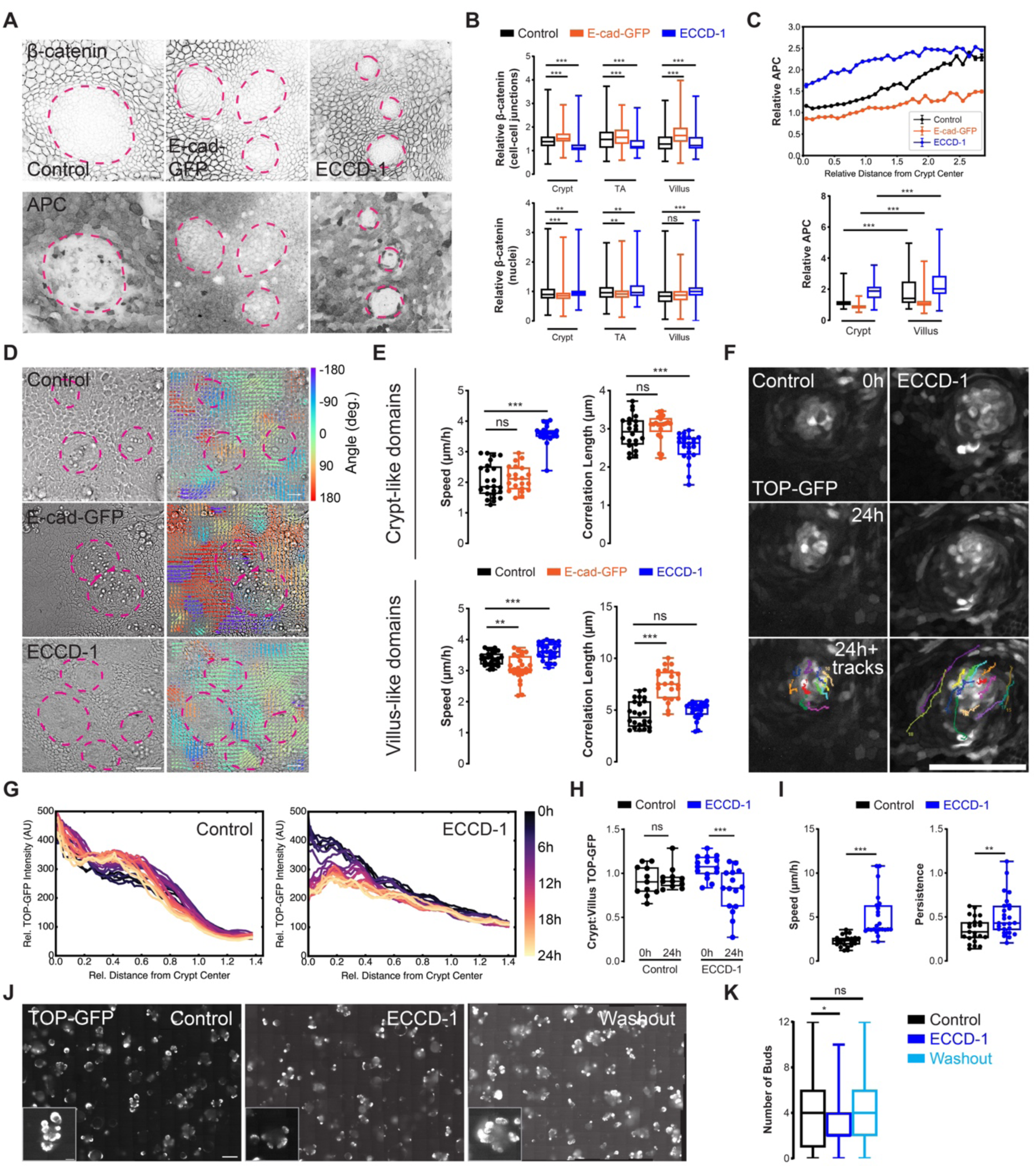
Modulation of cell-cell junctions dysregulates Wnt signaling and homeostasis via β-catenin. **A.** Representative micrographs of monolayers generated from untreated parental mTmG (control) organoids, mTmG/E-cad-GFP (Cdh1-GFP) overexpressing organoids and wild type organoids treated with a functional antibody targeting the extracellular domain of E-cadherin (ECCD-1) and immunostained for β-catenin (single z-slice) and APC (3D projection). Magenta dashed lines indicate crypt-like regions. SB: 50 μm. **B.** Quantification of relative β-catenin and APC fluorescence intensity in subcellular compartments (cell-cell junctions, nuclei) of individual cells from crypt, transit amplifying (TA) and villus compartments for organoid monolayers as in *A*. Intensity of protein at cell-cell junctions and nuclei are relative to cytoplasmic intensity from individual cell segmentations. Data compiled from *n* = 15, 16, 15 monolayers from *N* = 3, 3, 3 independent experiments. **C.** *Top*, Binned linescans of relative APC intensity along a computationally reconstructed crypt-villus axis for conditions in *C*. The relative intensity of each cellular compartment is normalized with cytoplasmic signal. Position values along the crypt-villus axis are rescaled such that the crypt center is at position × = 0, the crypt edge at position × = 1, and the villus-like compartment at values × > 1. Binned intensity values indicate mean ± SEM. *Bottom*, boxplots of relative intensity of APC in different intracellular compartments of individual cells from crypt, TA and villus compartments. Data compiled from same immunostainings as shown in *B*. **D.** Representative brightfield micrographs and particle image velocity (PIV) analysis of monolayers generated from control organoids, organoids stably over-expressing E-cadherin or organoids treated with ECCD-1. SB: 50 μm. Scale vectors: 12 μm/hr. **E.** Quantification of mean instantaneous speeds and correlation lengths for crypt-like and villus-like domains of organoid monolayers analyzed by PIV as in *D.* Each dot represents one organoid monolayer from *n* = 20, 22,21 monolayers and *N* = 3, 3 independent experiments. **F.** Representative micrographs of TOP-GFP organoid monolayers with or without ECCD-1 treated. Individual TOP-GFP^hi^ cells tracked by live microscopy for 24 hours (colored tracks). SB: 50 μm. **G.** Radially averaged intensity linescans from the center of crypt-like region (x = 0) to the villus like region over time from the examples shown in *F*. **H.** Quantification of change in total GFP intensity for TOP-GFP^hi^ cells in crypt-like and villus-like compartments expressed as a ratio of the final timepoint (t = 24 h) to the initial timepoint (t = 0 h) as for treatments in *F*. Each dot represents one organoid monolayer from *n* = 21, 20 and *N* = 3, 3 independent experiments. **I**. Quantification of migration dynamics of individual TOP-GFP^hi^ cells for treatments as in *F*. Each dot represents an individual TOP-GFP^hi^ cell from *n* = 7, 7 organoid monolayers and *N* = 3, 3 independent experiments. **J.** Representative micrographs of 3D TOP-GFP organoids treated with ECCD-1 and after inhibitor washout (72 h). SB: 200 μm; inset: 50 μm. **K.** Quantification of organoid budding for conditions as in *J*. Data compiled from *n* = 52, 50, 55 organoids from *N* = 3, 3, 3 independent experiments. Statistical tests performed using one-way ANOVA tests (p<0.001 for all plots) and Bonferroni’s post-hoc test (*p<0.05, **p<0.01, ***p<0.001, ns:p>0.05).

Our previous results suggest that modulation of Wnt signaling influences tissue compartmentalization and migration dynamics. To investigate how directly interfering with cell-cell junctions influences tissue dynamics, we performed PIV analysis of organoid monolayers with E-cadherin overexpression or blocking using ECCD-1 (Figure 3D; Video S4). While E-cadherin overexpression had no significant effect on migration speed or correlation length in crypt-like compartments, ECCD-1 treatment led to a significant increase in speed and a reduction in correlation length in both the crypt- and villus-like compartments, characteristic of tissue unjamming^40,46^ (Figure 3E). In the villus-like compartment of mTmG/E-cad-GFP monolayers, we observed a decrease in migration speed and an increase in correlation length, indicating tissue jamming.

To investigate how disruption of cell-cell junctions influences Wnt signaling and crypt dynamics on the single-cell level, we engineered an organoid line stably expressing TOP-GFP, a fluorescent reporter of active Wnt signaling^47^ (Figure S2C). We then generated TOP-GFP organoid monolayers, treated with ECCD-1 and performed live imaging (Figure 3F; Video S5). To analyze the spatial distribution of TOP-GFP over time, we generated intensity linescans from the center of the crypt-like region to the villus-like region over time, averaging the linescans radially around the entire crypt-like domain (Figure 3G). The resulting average TOP-GFP linescans in untreated organoids were stable over time, while ECCD-1 treated monolayers led to a loss of TOP-GFP intensity at the center of the crypt and increased migration dynamics of TOP-GFP^hi^ cells (Figure 3H,I). To determine whether disruption of cell-cell junctions influences crypt compartment specification, we quantified the budding of 3D organoids following treatment and/or washout of ECCD-1 (Figure 3J,K). ECCD-1 treatment led to reduced bud formation, which could be rescued by subsequent washout. Performing live imaging of TOP-GFP organoids revealed that ECCD-1 increased oscillatory migration of TOP-GFP^hi^ cells and reduced bud formation time (Figure S2D,E; Video S6). This suggests that increasing tissue fluidity by reducing cell-cell junction integrity allows for more efficient tissue rearrangement during bud specification, at the expense of the number of bud regions. Taken together, these data indicate that direct perturbation of cell-cell junctions can modulate Wnt signaling by β-catenin sequestration or release at junctions, and that this functionally interferes with crypt specification.

### Perturbation of Wnt signaling disrupts intestinal crypt specification and tissue dynamics

We next sought to determine whether the disruptions in collective dynamics and tissue organization observed following cell-cell junction modulation are consistent with phenotypes arising from direct Wnt pathway perturbation. To this end, we performed brightfield microscopy to probe changes in overall organoid morphology and growth following CHIR and LGK treatment and subsequent washout (Figure 4A). Organoid budding was severely perturbed following CHIR treatment but was restored following washout, while the loss of budding following LGK treatment was irreversible (Figure 4B). In addition, LGK treatment led to an increase in cell height along the apico-basal axis as well as a reduction in the roundness of the organoid lumen (Figure 4C; Figure S3A,B).

**Figure 4.**
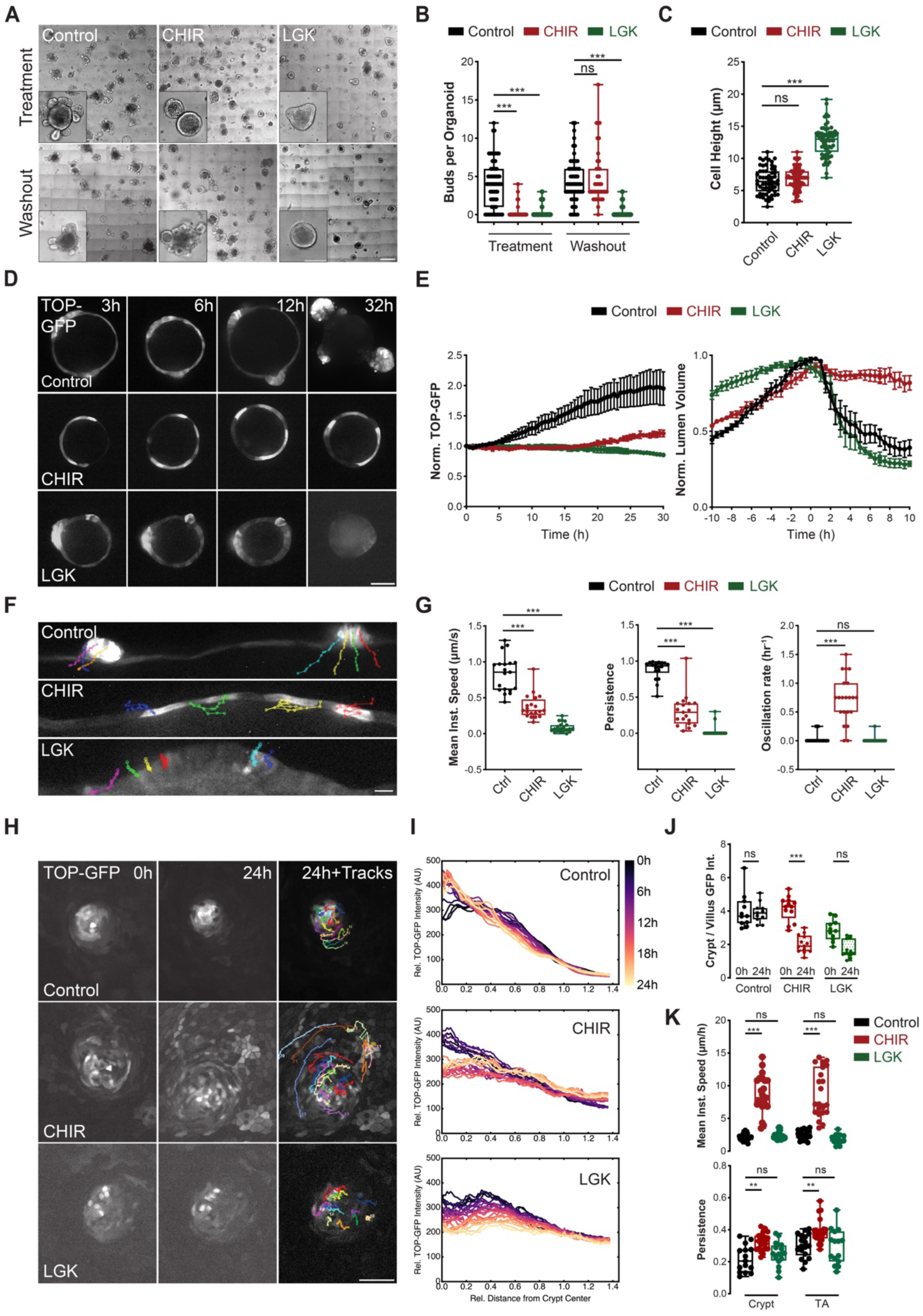
Perturbation of Wnt signaling disrupts intestinal crypt specification and stability. **A.** Representative brightfield micrographs of 3D intestinal organoids treated with CHIR or LGK and after inhibitor washout (72 h). Control organoids were untreated. SB: 200 μm, inset: 50 μm. **B.** Quantification of organoid budding for conditions as in *A*. Each dot represents one 3D organoid from *n* = 70, 75, 70, 60, 60,60 organoids and *N* = 4, 4, 4, 4, 4, 4 independent experiments. **C.** Quantification of cell height in 3D organoids following CHIR or LGK treatment as in *A*. Each dot represents one 3D organoid from *n* = 40, 40, 40 organoids and *N* = 4, 4, 4 independent experiments. **D.** Representative micrographs of 3D TOP-GFP organoids pre-treated with CHIR or LGK prior to live imaging. SB: 50 μm. **E.** Quantification of mean TOP-GFP expression in whole organoids (*left*) and organoid lumen volume (*right*) for conditions as in *D* over time. Data points reflect mean ± SD from *n* = 30, 32, 30 organoids and *N* = 3, 3, 3 independent experiments. **F.** Representative straightened images 3D TOP-GFP organoids from *D* with tracking of individual TOP-GFP^hi^ cells over time (colored tracks). SB: 20 μm. **G.** Quantification of mean instantaneous speed, persistence and directional oscillation rate of TOP-GFP^hi^ cells tracked as in *F*. Each dot represents one of *n* = 22, 25, 21 TOP-GFP^hi^ cells (6, 7, 7 organoids) from *N* = 3, 3, 3 independent experiments. **H.** Representative micrographs of TOP-GFP organoid monolayers pre-treated with CHIR or LGK prior to performing live confocal microscopy. Individual TOP-GFP^hi^ cells were tracked over time (colored tracks). SB: 50μm. **I.** Radially averaged intensity linescans from the center of crypt-like region (x = 0) to the villus like region over time from the examples shown in *H*. **J.** Quantification of change in total GFP intensity for TOP-GFP^hi^ cells in different compartments (crypt, transit amplifying [TA], villus) expressed as a ratio of final timepoint (t = 29h) to initial timepoint (t = 0 h) as for treatments in *H*. Each dot represents one crypt-villus structure from *n* = 6, 6, 6 organoid monolayers and *N* = 3, 3, 3 independent experiments. **K.** Quantification of migration dynamics of individual TOP-GFP^hi^ cells in crypt- or TA-like regions after treatments as in *H*. Each dot represents one TOP-GFP^hi^ cell from *n* = 25, 26 cells and *N* = 3, 3 independent experiments. Statistical tests performed using one-way ANOVA tests (p<0.001 for all plots) and Bonferroni’s post-hoc test (**p<0.01, ***p<0.001, ns:p>0.05).

To more directly investigate how disruption of the Wnt pathway influences downstream signaling dynamics and stem cell organization, we performed live confocal imaging of TOP-GFP organoids following CHIR or LGK treatment using freshly passaged organoids prior to budding (Figure 4D; Video S7). In control organoids, all cells expressed a moderate baseline level of TOP-GFP, with several highly expressing cells (TOP-GFP^hi^) in discrete regions of the organoid. Within ∼24 hours, buds formed at sites corresponding to the regions containing TOP-GFP^hi^ cells, while TOP-GFP expression in the neighboring cells progressively decreased. Following bud formation, TOP-GFP expression increased significantly at the base of the mature buds. In CHIR-treated organoids, we observed a similar initial heterogeneity in TOP-GFP expression, with distinct TOP-GFP^hi^ cells. However, the remaining cells did not lose TOP-GFP expression over time, and no buds were formed. In LGK-treated organoids, regions containing TOP-GFP^hi^ cells appeared to initially specify buds, which began protruding. However, over time, all cells displayed a reduction in TOP-GFP expression, and the nascent bud regions eventually regressed. Quantifying mean TOP-GFP expression over time, we found that in control organoids, expression gradually increased over time before reaching a plateau after ∼25 hours (Figure 4E). In contrast, CHIR-treated organoids displayed only a minor increase in overall TOP-GFP expression starting at ∼20 hours, while TOP-GFP expression gradually decreased over time in LGK-treated organoids. Measuring organoid lumen volume over time with respect to initial bud formation, we found that control organoids underwent an initial swelling prior to budding followed by compaction as the bud “neck” region closed, consistent with previous reports^48^. CHIR-treated organoids also initially swelled and reached a plateau in volume without budding, while the lumens of LGK-treated organoids initially swelled before gradually shrinking over time.

To determine how differing cell dynamics contribute to bud formation and collapse under these conditions, we analytically unwrapped the organoids along the equator and tracked individual cell movements (Figure 4F; Video S8). In control organoids, TOP-GFP^hi^ cells migrated in a convergent manner during bud specification, coinciding with the characteristic bending associated with budding morphogenesis. In CHIR-treated organoids, TOP-GFP^hi^ cells displayed a ∼2-fold reduction in migration speed, with low migration persistence and periodic oscillations, leading to the maintenance of a constant spacing between TOP-GFP^hi^ cells over time (Figure 4G). TOP-GFP^hi^ cells in LGK-treated organoids displayed severely reduced migration dynamics.

To better understand how these differences in the dynamics of TOP-GFP^hi^ cells influence crypt organization, we generated organoid monolayers using TOP-GFP organoids, treated with CHIR or LGK and measured the change in crypt area over time (Figure S3C,D). While control monolayers remained relatively stable, with a ∼10% decrease in crypt area over ∼20 hours, CHIR-treated monolayers displayed a ∼50% increase in crypt area. LGK-treated monolayers, on the other hand, displayed a ∼50% decrease in crypt area over ∼20 hours. To better understand how disruption of Wnt signaling interferes with cell dynamics and signaling within the crypt compartment, we tracked TOP-GFP reporter expression and individual cell motility over time (Figure 4H; Video S9). Following CHIR treatment, TOP-GFP intensity at the center of the crypt-like region gradually decreased, while intensity concomitantly increased in the villus like region, leading to a dissolution of the Wnt signaling gradient over time, similar to the effect of ECCD-1 treatment (Figure 4I). Following LGK treatment, TOP-GFP decreased in the crypt-like domain over time, leading to a flat, low profile, consistent with the loss of Wnt signaling. Comparing TOP-GFP intensity in the crypt-like and villus-like compartments, we found that while control organoids maintained a high crypt:villus TOP-GFP ratio, this ratio decreased to ∼1 in CHIR- and LGK-treated organoids, indicating a loss of compartmentalization (Figure 4J). TOP-GFP^hi^ cells demonstrated a ∼5-fold increase in migration speed and a ∼1.5-fold increase in migration persistence following CHIR treatment, while LGK treatment did not significantly change migration speed or persistence (Figure 4K). Together, these results suggest that over-activation of Wnt signaling by GSK3β inhibition leads to the expansion of Wnt signaling competent progenitors from crypts, which are unable to undergo budding morphogenesis. Repression of Wnt signaling by LGK treatment, conversely, leads to a gradual loss of stem cells with high Wnt signaling and organoid death within ∼2-3 days, likely due to spontaneous differentiation and the limited life span of differentiated intestinal epithelial cells.

### Wnt signaling regulates intestinal stem cell differentiation and tissue compartmentalization

The loss of TOP-GFP^hi^ cells and reduced organoid viability following Wnt repression suggests that in the absence of Wnt, intestinal stem cells spontaneously differentiate. To test this, we treated organoids with CHIR or LGK before fixing and immunostaining for the cell proliferation marker Ki-67 and Olfactomedin 4 (OLFM4), a marker of adult intestinal stem cells (Figure 5A). While OLFM4 and Ki-67 expression was restricted to the crypt-like bud regions in control organoids, cells in CHIR-treated organoids displayed homogeneous OLFM4 and Ki-67 expression throughout the organoid. In LGK-treated organoids, OLFM4 expression was severely reduced, and Ki-67 expression was restricted to a small fraction of cells at nascent buds (Figure 5B). These data suggest indicate that perturbation of upstream Wnt signaling regulates intestinal stem cell differentiation. This is also consistent with the dysregulation of budding following treatment with CHIR or LGK, as symmetry breaking during budding has been linked with local signaling heterogeneities and differentiation^49,50^.

**Figure 5.**
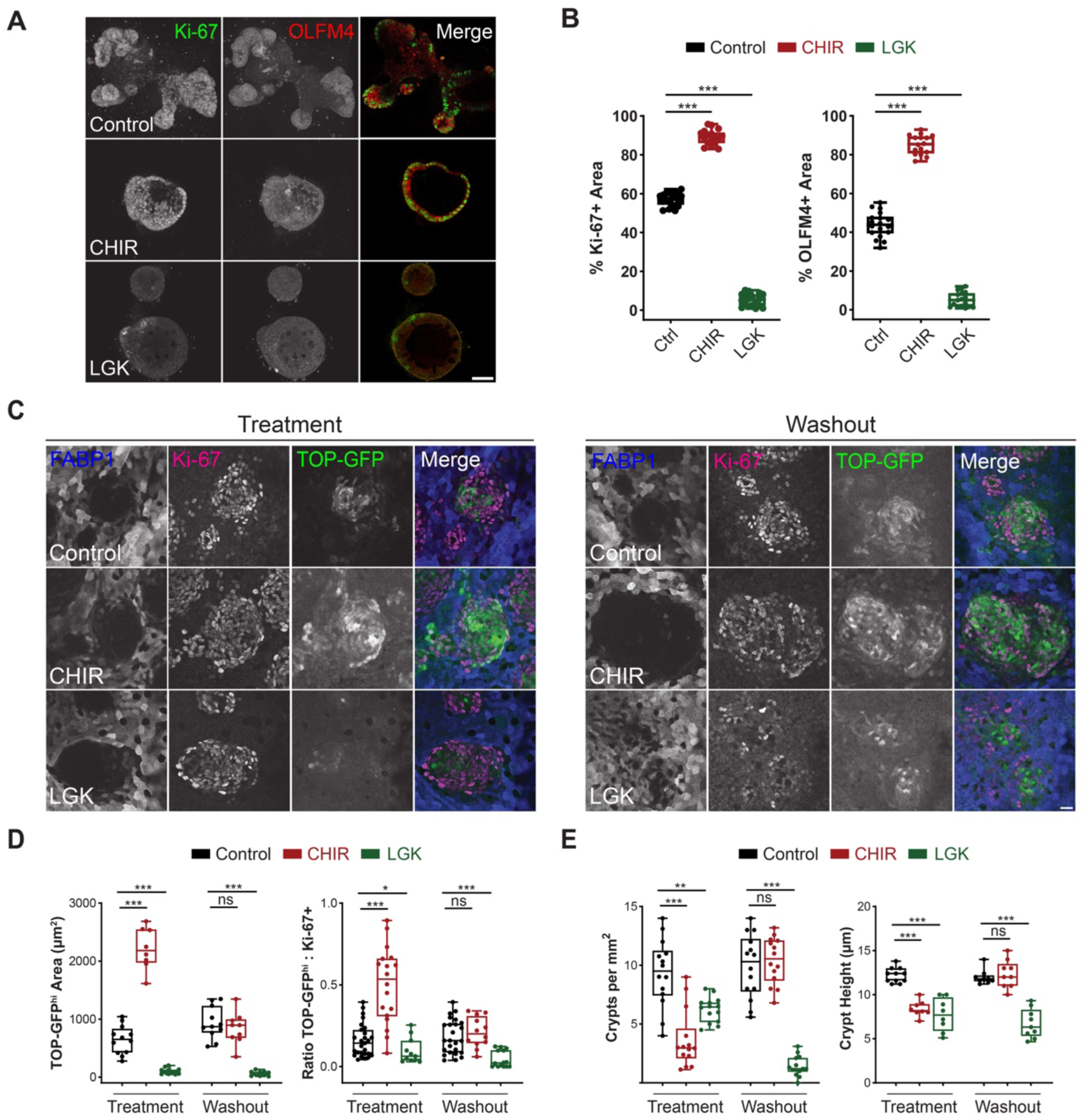
Wnt signaling regulates intestinal stem cell organization and differentiation in organoids. **A.** Representative micrographs of 3D organoids treated with treated with CHIR or LGK and immunostained for Ki-67 and Olfactomedin 4 (OLFM4). Images are 3D-projections of z-stacks. SB: 50µm. **B.** Quantification of percentage of Ki-67 and OLFM4 positive area relative to total organoid area for conditions in *A*. Each dot represents one 3D organoid from *n* = 20, 24, 19, 20, 19, 20 organoids and *N* = 3, 3, 3, 3, 3, 3 independent experiments. **C.** Representative micrographs of TOP-GFP organoid monolayers treated with CHIR or LGK with or without treatment washout and immunostained for fatty acid-binding protein 1 (FABP1) and Ki-67. SB: 20 μm. **D.** Quantification of absolute and relative TOP-GFP area with respect to the Ki-67 positive area as for conditions as in *C*. **E.** Quantification of the frequency and height of crypt-like compartments for conditions as in *C*. For *D* and *E*, each dot represents one crypt-villus structure from *n* = 6, 6, 7 organoid monolayers and *N* = 3, 3, 3 independent experiments. Statistical tests performed using one-way ANOVA tests (p<0.001 for all plots) and Bonferroni’s post-hoc test (*p<0.05, ***p<0.001, ns:p>0.05).

To further investigate how perturbation of Wnt signaling influences intestinal stem cell differentiation, we treated TOP-GFP organoid monolayers with CHIR or LGK and performed subsequent inhibitor washout, fixed and immunostained organoids for fatty acid-binding protein 1 (FABP1) to label differentiated enterocytes and Ki-67 to label proliferating cells (Figure 5C). Although both inhibitors are reversible, differentiation itself is effectively irreversible under homeostatic conditions. We therefore expected that stimulation of Wnt signaling by CHIR would drive a reversible increase in stem-cell like behavior, while even a temporary blockade of Wnt signaling by LGK would lead to an irreversible loss of stem cells. In line with this prediction, control organoid monolayers displayed robust compartmentalization of Ki-67-expressing crypt-like domains and FABP1-expressing villus-like domains. In CHIR-treated monolayers, the contours of the crypt-like domains appeared more convoluted, indicating reduced compartmentalization. LGK-treated monolayers initially maintained well-organized boundaries between crypt- and villus-like domains with low levels of TOP-GFP expression. Following washout, normal compartmentalization was restored in CHIR-treated organoids, while in LGK-treated organoids, the crypt domains gradually disappeared due to spontaneous differentiation of stem and progenitor cells, resulting in a more homogeneous population of FABP1-expressing cells. Quantification of the size of the TOP-GFP^hi^ domain confirmed that CHIR treatment led to a reversible expansion of the central crypt-like domain, while LGK treatment led to an irreversible loss of the TOP-GFP^hi^ domain (Figure 5D). Consistent with this, the frequency of crypt-like domains was significantly reduced following CHIR treatment but recovered following washout, while crypt-like regions disappeared over time in LGK-treated monolayers, even after washout (Figure 5E). Both CHIR and LGK treatment also led to a significant decrease in cell height in the remaining crypt-like regions, consistent with previous work showing that cells in the crypt-like compartments are more columnar due to reorganization of the actomyosin cytoskeleton^35^. Following washout, crypt-like regions of CHIR-treated monolayers were restored and recovered their normal height, while crypt-like regions in LGK-treated monolayers did not recover and maintained a reduced height. Together, these data confirm that Wnt signaling is a central regulator of intestinal stem cell differentiation and maintenance of boundaries between crypt- and villus-like domains.

### Modulation of upstream Wnt signaling perturbs tissue compartmentalization and localization of destruction complex components

Our results suggest that perturbation of Wnt signaling leads to changes in the intracellular localization and dynamics of β-catenin as well as modifications in downstream signaling activity. To better understand the link between β-catenin localization and activation of Wnt transcriptional targets in the context of canonical Wnt signaling, we fixed TOP-GFP organoid monolayers and immunostained for β-catenin and the destruction complex component APC (Figure S3E). TOP-GFP^hi^ cells formed a sharp interface at the edge of the crypt-like compartment with cells expressing low levels of TOP-GFP (TOP-GFP^low^). TOP-GFP expression was positively correlated with nuclear, but not junction-associated, β-catenin (Figure S3F). The boundary between TOP-GFP^hi^ and TOP-GFP^low^ cells also delineated APC expression, with TOP-GFP^hi^ cells expressing low APC and TOP-GFP^low^ cells expressing high APC. To investigate how perturbation of Wnt signaling influences APC localization, we treated organoid monolayers with CHIR or LGK and immunostained for β-catenin and APC (Figure S3G). Treatment with either CHIR or LGK resulted in an increase in APC levels in crypt-like regions, in addition to an accumulation of APC at the apical membrane (Figure S3H). These data suggest that in addition to modulating β-catenin localization and dynamics, perturbation of Wnt signaling leads to changes in destruction complex localization.

### Constitutive activation of Wnt signaling by APC deletion disrupts crypt/villus compartmentalization, tissue dynamics and barrier function

While pharmacological perturbation experiments reveal how acute disruption of Wnt signaling impacts β-catenin organization and tissue dynamics, it is unclear how these effects reflect changes associated with tumorigenesis. To address this, we employed CRISPR/Cas-mediated genome editing to generate an APC knock-out (APC^−/-^) organoids (Figure 6A,B). Similar to CHIR-treated organoids, APC^−/-^ organoids lacked bud regions and displayed significantly reduced cell height (Figure 6C). Immunostaining of 3D organoids indicated that APC^−/-^ organoids primarily contained actively dividing Ki-67+ cells and lacked differentiated FABP1+ enterocytes (Figure 6D,E). To validate that APC^−/-^ organoids displayed similar behavior to intestinal tumors, we used tumor organoids (tumoroids) harvested from the inducible spontaneous NICD/p53^−/-^ mouse model^51^. Similar to APC^−/-^ organoids, NICD/p53^−/-^tumoroids lacked distinct buds, and APC could not be detected by Western blot or immunofluorescence (Figure S4A-D). Mouse tumoroids also contained mainly actively dividing Ki-67+ cells and mostly lacked differentiated FABP1+ enterocytes (Figure S4E,F).

**Figure 6.**
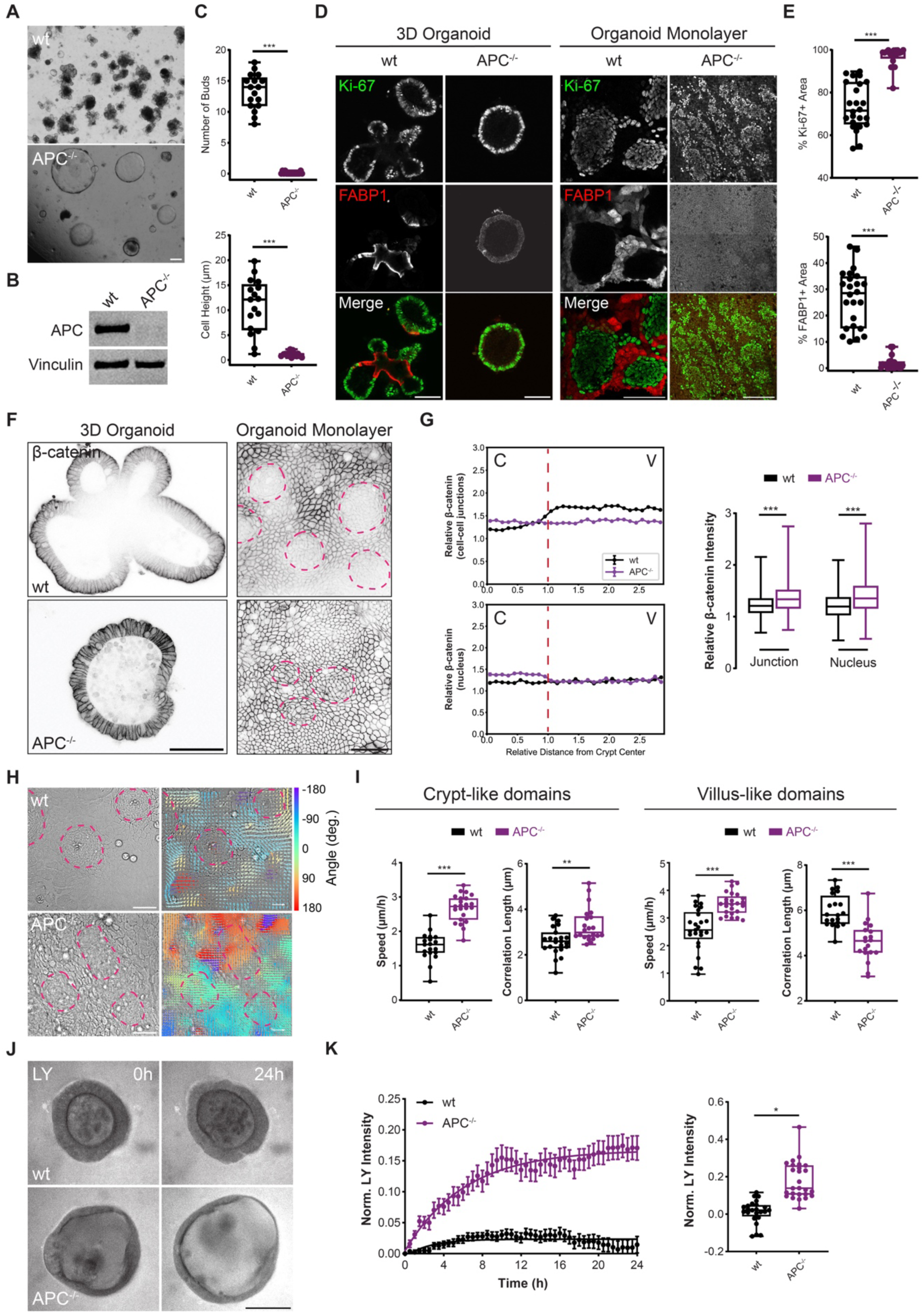
Constitutive activation of Wnt signaling by APC deletion disrupts crypt/villus organization, tissue dynamics and barrier function. **A.** Representative brightfield micrographs of 3D wild-type (wt) and APC knock-out (APC^−/-^) mouse intestinal organoids 7 days after passage. SB: 50µm. **B.** Western blot of APC expression in wt and APC^−/-^ organoids. Vinculin is used as a loading control. **C.** Quantification of number of buds and cell height from conditions in *A* from n=18,18 organoids from N=3,3 independent experiments. **D.** Representative micrographs of wt and APC^−/-^ organoids in 3D culture or monolayer culture immunostained for Ki-67 and FABP1. SB: 50 µm, 50 µm, 50 µm, 50 µm. **E.** Quantification of Ki-67 and FABP1 positive area relative to total monolayer area for conditions as in *D*. Each dot represents one organoid monolayer from *n* = 25, 22 monolayers and *N* = 3, 3 independent experiments. **F.** Representative micrographs of wt and APC^−/-^ in 3D culture or monolayer culture immunostained for β-catenin. Magenta dashed lines indicate crypt-like regions. SB: 50µm, 50 μm. G. *Left*, Binned linescans of relative β-catenin intensity in different intracellular compartments along a computationally reconstructed crypt-villus axis as for organoid monolayers under conditions in *F*. The relative intensity of is normalized with cytoplasmic signal. Position values along the crypt-villus axis are rescaled such that the crypt center is at position × = 0, the crypt edge at position × = 1, and the villus-like compartment at values × > 1. Binned intensity values indicate mean ± SEM. *Right*, boxplots of relative intensity of β-catenin of individual cells from crypt-like compartment. Data compiled from *n* = 16, 16, 16 organoids monolayers and *N* = 3, 3, 3 independent experiments. **H.** Representative brightfield micrographs and particle image velocity (PIV) maps of wt and APC^−/-^ organoid monolayers. SB: 50 µm. Scale vector: 12 μm/hr. **I.** Quantification of mean instantaneous speeds and correlation lengths for crypt-like and villus-like domains of organoid monolayers analyzed by PIV as in *H.* Each dot represents one organoid monolayer from *n* = 19, 23 monolayers and *N* = 3, 3 independent experiments. **J.** Representative micrographs of 3D intestinal organoids treated with CHIR or LGK, after adding Lucifer Yellow (LY) to the organoid medium. SB: 50µm. K. *Left*, Quantification of relative luminal fluorescence (organoid lumen to medium fluorescence intensity) over time after adding LY (mean ± SEM). *Right*, quantification of normalized luminal LY intensity after 21 h for condition as in *E*. Each dot in the box plot represents one organoid from *n* = 46, 45 and *N* = 3, 3 independent experiments. Statistical tests performed using one-way ANOVA tests (p<0.001 for all plots) and Bonferroni’s post-hoc test (*p<0.05, **p<0.01, ***p<0.001).

To determine how APC deletion influences intracellular β-catenin localization, we immunostained APC^−/-^ organoids in 3D and monolayer culture formats for β-catenin (Figure 6F). Quantifying subcellular localization of β-catenin along the spatially reconstructed crypt-villus axis in organoid monolayers revealed that APC^−/-^ organoids lacked a β-catenin gradient and displayed a slightly higher relative nuclear β-catenin localization compared with wt organoids (Figure 6G). We validated this change in intracellular localization using biochemical fractionation, which indicated that APC^−/-^ organoids had higher overall expression of β-catenin as well as increased nuclear β-catenin, but decreased cytoplasmic β-catenin (Figure S5A,B). APC^−/-^ organoid monolayers also displayed increased localization of NUP98 to the nuclear cortex (Figure S5C,D). Taken together, these findings are consistent with the reduction in the β-catenin gradient along the crypt-villus axis and increase in nuclear β-catenin import following activation of Wnt signaling via CHIR treatment.

As activation of Wnt signaling following CHIR treatment promoted collective cell migration and increased tissue fluidity (Figure 2), we next investigated whether APC deletion also led to dysregulation of migration dynamics. To this end, we performed PIV analysis on wt and APC^−/-^ organoid monolayers (Figure 6H; Video S10). APC^−/-^ organoids displayed increased migration velocity and correlation length in the crypt-like domain, similar to CHIR treatment, and decreased correlation length in the villus-like domain (Figure 6I). As increased tissue fluidity is likely associated with reduced cell-cell junction integrity, we next probed barrier function in APC^−/-^ organoids using the live imaging-based LY assay (Figure 6J; Video S11). We observed that APC^−/-^ organoids indeed displayed reduced barrier function, indicated by increased LY permeation into the organoid lumen (Figure 6K). Together, these data confirm that disruption of the destruction complex by either pharmacological or genetic perturbation led to aberrantly increased Wnt signaling as well as dysregulation of normal tissue dynamics and organization. This raises the possibility that early stages of tumorigenesis, which are commonly driven by APC mutations in the intestine, may also perturb intestinal barrier integrity, which could lead to increased local inflammation, further driving tumor progression.

## Discussion

Here, we investigated the interplay between the signaling and adhesion functions of β-catenin in the small intestinal epithelium, which is highly dependent on Wnt signaling for homeostasis and regeneration. Our data suggest that driving Wnt signaling either by pharmacological or genetic abrogation of destruction complex components induced changes in intracellular β-catenin localization and dynamics and led to tissue fluidification and loss of epithelial integrity, suggestive of disrupted cell-cell adhesion. Direct stabilization of cell-cell junctions by E-cadherin overexpression led to recruitment of β-catenin at junctions at the expense of nuclear localization, while blocking E-cadherin binding led to β-catenin release from junctions, increased nuclear localization and higher Wnt signaling. Together, our findings indicate that the intracellular localization of β-catenin, in particular its association with cell-cell junctions tunes its downstream signaling function. This has major implications for understanding intestinal epithelial stem cell behavior during homeostasis and tumorigenesis.

β-catenin was discovered independently for its role at cell-cell junctions as a binding partner of E-cadherin and for its essential function in development^52–55^. Early structural analysis revealed that separate domains mediate the signaling and adhesion roles^56,57^. Studies in developing epithelial tissues in *Xenopus* and *Drosophila* raised the possibility of competition between the signaling and adhesions functions of β-catenin^58–63^. Later studies using cell lines suggested that while interfering with E-cadherin can lead to the release of β-catenin from junctions, this is not sufficient to promote downstream signaling^64–70^. Thus, these previous studies present conflicting models regarding the potential competition or independence of β-catenin’s role in Wnt signaling and cell-cell adhesions. While there is likely to be a component of cell-type specificity, our data can be interpreted as being consistent with both models. Our biochemical fractionation data indicate that while inhibition of GSK-3β by CHIR treatment led to a marked increase in cytoplasmic levels of β-catenin, nuclear levels were largely unchanged (Figure S1B,C). This suggests that, as expected from the commonly accepted model of canonical Wnt signaling^3,71^, perturbation of the disruption complex leads to excess β-catenin levels. However, as nuclear import is an active process, this may not directly translate to increased nuclear β-catenin. Furthermore, our biochemical analysis does not account for differences between cell types and tissue compartments. Our immunostaining data indicates that GSK-3β inhibition leads to a global increase in junction-associated β-catenin (Figure 1C,D). This treatment leads to increased nuclear β-catenin specifically in the crypt-like domain and loss of nuclear β-catenin in the villus-like compartment. The difference in this response suggests that certain cells, such as intestinal epithelial stem cells, are primed to respond to Wnt signaling. Consistent with this, in Apc^−/-^ organoid monolayers, we observed a flat profile of junctional β-catenin along the reconstructed crypt-villus axis, but an increase in nuclear β-catenin specifically in crypt-like compartments (Figure 6F,G). We additionally found that increased Wnt signaling by pharmacological or genetic perturbation promoted expression and accumulation of the nuclear pore protein NUP98 at the nuclear cortex, potentially driving further nuclear β-catenin import in the crypt. While our immunostaining experiments suggest changes in relative nuclear β-catenin localization, the accumulation of nuclear β-catenin is relatively subtle compared with nuclear β-catenin staining *in vivo* in intestinal stem cells and invasive colorectal cancer cells, which display both nuclear and junctional β-catenin^72,73^. Nevertheless, we are still able to measure a shallow gradient of nuclear β-catenin enrichment in the stem-cell rich crypt-like domain (Figure 1C,D). These data are also consistent with our biochemical fractionation results, where we primarily observe changes in cytoplasmic β-catenin levels (Figure S1B,C). This biochemical data, which lacks spatial information, also over-represents differentiated enterocytes, which make up a significantly larger fraction of the total cells compared to the stem cells.

Mechanical stresses at cell-cell adhesions can regulate cell proliferation, differentiation, tissue organization and barrier integrity^74–78^. Previous studies have identified various nuclear signaling proteins that localize to cell-cell junction complexes and whose localization depends on tissue integrity and junction maturation^79–87^. As the assembly of cell-cell junction complexes is known to be mechanosensitive^88–91^, junctional integrity itself could also trigger mechanotransduction through the release or sequestration of various signaling factors. Our experiments modulating E-cadherin indicate that β-catenin localization and activity can be directly influenced by junction stability. In addition to its sequestering function, E-cadherin has also been shown to increase β-catenin stability and may aid in buffering β-catenin concentrations and activity^14^. Phosphorylation of β-catenin has been suggested to add an additional regulatory level on top of localization and sequestration, potentially leading to distinct pools of β-catenin with different binding capacities and functions^11,13^. However, the expression of endogenous full length β-catenin, especially in cell types that require Wnt signaling like the intestinal epithelium, is still likely to result in a competition between β-catenin localization, post-translational modification and function. Indeed, our biochemical data do not indicate a major effect of β-catenin phosphorylation status on subcellular compartmentalization, though untreated controls appear to have higher levels of β-catenin in the nucleus with activating phosphorylations (Figure S1B,C). Future studies investigating the role of specific phosphorylation sites in nuclear β-catenin import will shed more light on the interplay between the signaling and adhesions functions.

Although genetic perturbation by knock-out of APC better reflects natural intestinal tumor progression, pharmacological perturbation of Wnt signaling allows to test more immediate downstream consequences. The rapid change in migration dynamics following CHIR treatment (Figure 2A,B) suggests that these effects may not depend on modulation of differentiation via altered Wnt signaling, which is expected to be affected after several days, but rather on protein-protein interactions. It is therefore likely that this change in tissue dynamics can be attributed to the direct destabilization of cell-cell junctions. Indeed, other studies indicate that disruption of cell-cell junctions leads to increased tissue fluidity^32–34^. Consistent with this, we observe a significant decrease in barrier function following CHIR treatment (Figure 2E,F). We also observe a loss of barrier integrity in APC^−/-^ organoids (Figure 6J,K). Interestingly, this could serve as a link between inflammation and tumor progression, as reduced barrier function during chronic intestinal inflammation increases the susceptibility to infection and sepsis^92^. Recent studies highlight that loss of barrier integrity (“leaky gut”) is associated with colorectal cancer in humans and that bacteria can invade tumors from the intestinal tract^93–96^. Moreover, previous reports demonstrate that the mouse colorectal cancer model APC^Min/+^ is associated with chronic inflammation and loss of barrier integrity^97^. In addition to effects on the microbiome, long-term disruptions in APC function leading to changes in differentiation profiles may therefore also contribute to reduced barrier function. Previous work also indicates that differentiation is concomitant with changes in expression of junctional proteins and membrane transporters^48^. While many studies propose that the link between colorectal cancer and inflammation is mediated by changes the microbiome (“dysbiosis”)^98–100^, our study suggests that there may also be a more direct mechanism involving the disruption of cell-cell junctions due to APC loss-of-function. As local inflammation can drive the stromal reaction, this points toward a potential feedback loop wherein early tumor mutations dysregulate barrier function, leading to chronic infection and inflammation which in turn promote tumor progression.

Our data indicate that perturbation of upstream Wnt signaling components not only interferes with β-catenin localization and expression, but also with components of the destruction complex. APC expression is highest outside of the crypt-like compartment and in TOP-GFP^low^ cells (Figure S3E,F), indicating a negative correlation between Wnt signaling and APC expression. This follows logically, as APC is required for repressing Wnt signaling in the absence of Wnt ligand. This further suggests that active Wnt signaling in the crypt during homeostasis may not only arise from protein-protein interactions that interfere with the activity of the destruction complex, but may also direct APC degradation, which could further potentiate Wnt signaling. However, this mechanism is likely to be more complex, as either increasing Wnt signaling by CHIR treatment or repressing Wnt signaling by LGK treatment both result in increased APC expression (Figure S3G,H). The observation that APC accumulates strongly to the apical membrane in the absence of Wnt ligand (following LGK treatment) is consistent with previous findings that APC accumulates at the apical membrane in polarized epithelial cells^101^. Our findings build on this, demonstrating that this apically associated pool of APC is regulated by Wnt signaling. Other studies suggest that upstream Wnt signaling can indeed regulate destruction complex localization in an APC-dependent manner^102^. However, these experiments rely on individual unpolarized cells, meaning that the specific localization changes may not be directly applicable to polarized intestinal epithelial cells.

While colorectal cancer is associated with mutations in various components of the Wnt pathway, APC truncation mutations are by far the dominant driver of intestinal tumorigenesis^103–105^. Our results suggest that the regulation of expression and intracellular localization of both β-catenin and APC may be more complex than is commonly considered for canonical Wnt signaling. Indeed, a recent report suggests that APC loss of function can activate additional signaling pathways independent of β-catenin, including Hippo/YAP signaling via stabilization of AJUBA^106^. Interestingly, AJUBA can also be regulated by changes in cytoskeletal tension^77^, potentially linking the observed differences in cell dynamics and cell shape associated with modulated Wnt signaling. Furthermore, APC is known to have a number of additional functions, including acting as a mediator of actin-microtubule interactions, actin polymerization and focal adhesion turnover^107–109^. Recent work has demonstrated that APC truncation mutants can also perturb plasma membrane cholesterol, modulating the formation of membrane microdomains that facilitate Wnt signaling^110^. Future studies focused on linking these diverse functions of APC and cross-talk between different signaling pathways will provide more insight to explain why APC is the dominant Wnt-related tumor driver in colorectal cancer.

Our study primarily uses mouse small intestinal organoids to study the regulation of β-catenin and Wnt signaling in homeostasis and tumorigenesis. While this could raise questions about the relevance to colorectal cancer in humans, spontaneous intestinal cancer mouse models, including the APC^Min/+^ and NICD/p53^−/-^ models, develop tumors primarily in the distal small intestine rather than the colon^51,111^. While this site-specificity differs between mice and humans, this suggests that the small intestine is the relevant mouse organ to use for modeling tumorigenesis. Though the mechanisms underlying preferential sites of tumor formation and the differences between humans and mice are yet not well understood, this presents interesting avenues of research for future investigation.

Taken together, our study indicates that Wnt signaling during intestinal homeostasis and tumorigenesis depends not only on the regulation of β-catenin expression and cytoplasmic concentrations, but also on intracellular localization and competition for β-catenin’s adhesion and signaling functions. We propose that the current understanding of canonical Wnt signaling in the intestinal epithelium should be extended to incorporate these additional elements. A more comprehensive understanding of this pathway is essential for gaining deeper insight into intestinal pathophysiology.

## Data availability

Primary data will be shared by the lead contact upon reasonable request. Custom code written for 3D segmentation as well as the StarDist segmentation model will be made available open-source via github upon publication. Other custom code for image and data analysis will be shared by the lead contact upon reasonable request.

## Acknowledgments

We thank the members of the Clark lab for insightful discussions and critical reading of the manuscript. We also thank François Fagotto (CRBM), Danijela Matic Vignjevic, René-Marc Mège, Kerstin Feistel, Valentina Trivigno, Eliane Klein and Elisabeth Letellier (Uni. Luxembourg) for helpful discussions and advice as well as feedback on the manuscript. The authors gratefully acknowledge the Technology Platform “Cellular Analytics” of the Stuttgart Research Center Systems Biology for their support and assistance in this work. We thank Jean-Léon Maître for the Cdh1-GFP mouse line and Reda Bouras and Danijela Matic Vignjevic for generating the Cdh1-GFP organoids and for providing the NICD/p53^−/-^ tumor organoids. We also acknowledge the staff of the animal facility of the Institute of Cell Biology and Immunology at the University of Stuttgart for their care of mouse stocks and support for experimental procedures. T.N., S.S. and A.G.C. were supported by the Federal Ministry of Education and Research (BMBF) and the Baden-Württemberg Ministry of Science (MWK-BW) as part of the Excellence Strategy of the German Federal and State Governments (NWG-GastroTumors to A.G.C.). This work also received support from the Terra Incognita Program from the University of Stuttgart.

## Author contributions

T.N. and A.G.C. designed the research and wrote the manuscript. T.N. carried out the experiments and image analysis. A.G.C. wrote and tested custom software for image and data analysis. F.G. performed and analyzed some of the 3D organoid experiments. S.S. provided assistance for some experiments. J.S. and P.R. designed and aided in generation of APC^−/-^organoids. D.T. performed some of the RT-qPCR experiments. A.H. supported the RT-qPCR experiments and generation of APC^−/-^ organoids. All authors discussed the results and manuscript.

## Declaration of competing interests

The authors declare no competing interests.

## Methods

### Reagents

All critical reagents used in experimental procedures are listed in the supplemental information (Table S1).

### Animals

To generate mouse intestinal organoids, wild type C57BL/6J or C57BL6/N (male or female) mice ranging from eight weeks to six months old were used. All wild type mice were originally sourced via Charles River Europe and housed under specific-pathogen-free conditions (12 h light/dark cycle, ad libitum access to food and water). Animals were euthanized by CO_2_ asphyxiation followed by cervical dislocation, and tissues were collected immediately post-mortem. mT/mG mice^112^ were purchased from JAX via Charles River Europe and imported in to the local mouse quarantine facility. mT/mG mice were housed for 1-2 weeks prior to sacrifice for the purpose of harvesting intestinal organs to generate organoids. All housing procedures and euthanasia were performed in accordance with local and regional laboratory animal regulations, and experiments were approved by the local ethics board (relevant project IDs approved by the University of Stuttgart ethics board: IIG§4_2_2021.Clark, IZI§4-3-2022, IIG§4_1_2023, IZI§4_3_2024).

### Mouse small intestinal crypt isolation

Isolation of small intestinal crypts was performed as in previous studies^113^. Immediately following sacrifice, the small intestine (SI) was removed by incising distal to the pylorus and proximal to the caecum. The mesentery was stripped, luminal contents flushed with ice-cold Ca²⁺/Mg²⁺-free PBS (PBS-0), and the SI was opened longitudinally. After washing twice with ice-cold PBS-0, 15 mins each at 4°C, the tissue was cut into 2-3 mm fragments and incubated in ice-cold Dissociation Solution (2 mM EDTA, 1 mM DTT, 10 mM HEPES, pH 7.4) for 30 min on a rocking platform at 4°C. Crypts were released by vigorous pipetting: the first two fractions (villus-enriched) were discarded, and the remaining fractions (crypt-enriched) were filtered (100 µm, then 70 µm) and pelleted by centrifugation (300xg, 5 min, 4°C).

### 3D organoid culture

Crypt pellets were resuspended 1:1 in Cultrex Basement Membrane Extract (BME) and 4°C Resuspension Buffer (DMEM/F12 + 2% Anti/Anti), and 50 µL domes were plated in 24-well plates. After polymerization at 37°C for 30 min, 350 µL of serum-free ENR medium (Advanced DMEM/F-12, 1x GlutaMAX, 10 mM HEPES, 1x Pen/Strep, 1x B27 minus vitamin A, 1x N-2, 50 ng mL^−1^ EGF, 100 ng mL^−1^ Noggin, 500 ng mL^−1^ R-spondin-1) was added. ENR Medium was refreshed every 2-3 days. Cultures were passaged every 4-5 days by mechanical dissociation and re-embedding in fresh BME (500xg, 5 min, 4°C).

### Preparation of organoid monolayers

Preparation of organoid monolayers was adapted from previous work^35^. Glass-bottom culture dishes (35 mm diameter) were pre-coated by incubating with 300 μl water and 150 μl silane APTMS for 15 minutes. After removing excess silane and washing twice with water, dishes were soaked in water for 10 minutes with agitation, then dried with compressed air. The dishes were subsequently incubated for 30 minutes with 0.5% glutaraldehyde in PBS. Following extensive washing and drying, dishes were stored dry at 4°C until use.

Poly-A-Acrylamide (PAA) gels with a nominal stiffness of 5 kPa were prepared on the pre-coated glass bottom dishes using an established and validated formula (7.46% Acrylamide, 0.044% Bis-Acrylamide, 0.5% Ammonium Persulfate, 0.05% N,N,N’,N’-tetramethylethylenediamine; final concentrations [v/v])^35,114^. Then, 16 μl of gel solution was applied onto glass-bottom dishes and gently overlaid with an 18 mm diameter round coverslip. The gels were allowed to polymerize for exactly 60 min at room temperature, after which the coverslips were removed in PBS using a razor blade and forceps. Gels were washed briefly with PBS and stored at 4°C in PBS for up to one week.

Polydimethylsiloxane (PDMS) stencils were fabricated by mixing the base and curing agent at a 10:1 mass ratio, degassed via centrifugation (3200 RPM, 2 min, RT), poured into plastic petri dishes to a height of 2mm, and cured at 80°C for at least 1 hour to overnight. The cured PDMS sheets were cut into ∼2 cm squares using a razor blade and perforated using a biopsy punch to generate 2 mm holes. Stencils were stored dry and cleaned with 70% ethanol before reuse. PDMS stencils were passivated by incubation in 2% (w/v) Pluronic F-127 solution in PBS for 1 hour at room temperature with occasional swirling, followed by briefly rinsing with 70% ethanol and sterile water. Afterwards, stencils were washed twice with PBS, dried with compressed air, and air-dried for an additional 15 minutes.

One day before plating, PAA gels were activated by applying 50 μl of 2 mg/ml Sulfo-SANPAH solution and UV-irradiated (365 nm) using a UV Bench Lamp (Analytik Jena, Jena, Germany) for 7.5 minutes. Sulfo-SANPAH solution was then removed, and gels were washed with 10 mM HEPES 3 times for 2 mins each followed by one 1x PBS wash for 2 mins on an orbital shaker, then air-dried briefly. Activated gels were used within 30 minutes. Passivated PDMS stencils were placed on the activated gels. 10 μl of cold ECM coating solution containing collagen-I (250 μg/ml) and laminin-1 (100 μg/ml) in 10 mM HEPES was pipetted onto the stencil holes and incubated overnight at 4°C.

Organoids cultured in 3D were passaged one day prior to plating. On the plating day, BME domes containing organoids were incubated in Organoid Harvesting Solution at 4°C for 1 hour with gentle shaking. Organoids were then gently dissociated from the BME dome, centrifuged (500xg, 5 min), washed twice with 1x PBS (centrifuged at 500xg, 5min) and resuspended in ENR + 10 uM Y-27632 (15 μl per gel). This suspension was then seeded onto the ECM-coated regions of the PAA gel. After 1-2 hours incubation at 37°C, 5% CO_2_, 950 μl of fresh ENR medium was carefully added. Organoid monolayers were used 2-4 days later.

### Production of Wnt-reporter-GFP (TOP) and mEos2-β-catenin lentiviral particles

Lenti-X 293T cells were seeded onto 175 cm^2^ culture flasks at 70% of confluence in DMEM + 10% FBS 24h prior to transfection using PEI. A total 25 µg plasmid DNA was mixed with 75 µg of polyethylenimine (PEI) in OPTI-MEM. Lentiviral supernatants were prepared by co-transfection using a 3:2:1 ratio of envelope vector (pVSV-G^115^) : packaging (pax2) : plasmid of interest. The DNA mixture was incubated with 1:1 PEI in 1 ml optiMEM for 15-20 minutes at room temperature. The mixture was dropwise to HEK-293T Lenti-X 293T cells following replacement with fresh culture media. The culture medium was replaced 12 h after transfection, and viral supernatant was collected 24 h, 48 h and 72 h later. The viral supernatants were cleared by low-speed centrifugation, filtered through a 0.45 μm syringe filter and stored at 4°C until further processing. Viral supernatants were concentrated 185-fold by ultracentrifugation at 100,000 × g for 1 h 45 min. Viral pellets were resuspended in ENR+Transdux infection reagent divided in 300 µl aliquots (3D transduction) or 10 µl aliquots (for monolayer transduction) and stored at -80°C.

TOP-GFP was a gift from Ramesh Shivdasani (Addgene plasmid #35489; http://n2t.net /addgene:35489; RRID: Addgene_35489). mEos2-β-Catenin-20 was a gift from Michael Davidson (Addgene plasmid #57350; http://n2t.net/addgene:57350; RRID: Addgene_57350). pCMV-VSV-G was a gift from Bob Weinberg (Addgene plasmid #8454; http://n2t.net/addgene:8454; RRID:Addgene_8454). psPAX2 was a gift from Didier Trono (Addgene plasmid #12260; http://n2t.net/addgene:12260; RRID:Addgene_12260).

### Lentiviral transduction in 3D organoids

For the transduction procedure, two 50 μl BME2 domes of highly concentrated organoids (∼500 organoids total) at day 4 after extraction were split and cultured during 3 days in ENR supplemented with 10 µM CHIR-99021, 10mM Nicotinamide and 10 µM Y-27632 dihydrochloride (ENR++). Organoids were harvested from BME, washed and pelleted as described above. The pellet was suspended in 250 µL of TrypLE dissociation reagent and incubated for 2.5 min at 37°C followed by gentle pipetting. Next, 5ml of ENR + 10 μM Y27632 + 10% FBS was added to the suspension, and the dissociated organoids were washed 2 times with 37°C ENR + 10 μM Y27632 by centrifugation (350 xg, 4 min). The pellet was suspended in 300 µl of TOP-lentiviral particles as described above. The suspension was pitetted into the well of a standard 48-well plate pre-coated with a 180 μl layer of polymerized BME2 and incubated overnight at 37°C with 10% CO_2_ and 100% humidity. The following day, the unattached cells were removed with warm ENR++. A second layer of BME was added, allowed to be polymerized during 60 min in the incubator and filled with ENR++ medium. At 48 hours after infection, the medium was replaced with normal ENR and renewed every 2-3 days. Infected organoids were cultured for a period of 4 months until multiple buds were observed. Stably positive GFP-TOP organoids were picked individually by hand using a transmitted fluorescence microscope, deposited into BME2 domes and amplified.

### Lentiviral transduction in organoid monolayers

Monolayer of organoids were prepared as describe above with organoids pellet resuspended in 2 µl of mEos2-β-Catenin viral particle for each sample. After that the organoids were attached, the stencil was peeled off and 1mL of ENR++ was gently added. Following overnight incubation at 37°C with 10% CO2 and 100% humidity, the medium was replaced with normal ENR and was replaced every day.

### Generation of APC-KO organoids

Organoids to be electroporated were cultured, stimulated and prepared as described above for organoids infected with lentivirus. APC-KO organoids were generated by delivering a Cas9 ribonucleoprotein (RNP) complex, containing an Alt-R CRISPR-Cas9 guide RNA (crRNA:tracrRNA duplex or sgRNA) and recombinant Cas9 into TrypLE-digested organoids using the Neon^TM^ Transfection System (Thermo Fisher Scientific) following the manufacturer’s protocol. Shortly, 1 µl of RNP complex was formed by mixing 0,5 µl Cas9 enzyme with 0,2 µl Buffer R (Neon kit) and 0.5 µl gAPC-RNA for 15 minutes at room temperature. TrypLE-digested organoids were suspended in 9 µl of Buffer R and 1 µl of RNP complex above. Cells were electroporated using Neon Transfection System with the following settings: 2000 V, 2 ms pulse width, 2 pulses. The electroporated cells were left untouched for 30 sec and then deposited into 48-well plates pre-coated with a layer of polymerized BME2 and covered with ENR++ medium. Cells were incubated overnight at 37°C with 10% CO2 and 100% humidity. The next day, unattached and dead cells were washed with ENR pre-warmed to 37°, and remaining attached cells were sandwiched with second layer of BME and 350 µl of ENR++ was added after polymerization of top layer of BME2. At 48 hours, the medium was switched to normal ENR and replaced every 2-3 days. After 3-4 weeks, APC KO organoids were selected based on their large, cystic appearance, manually picked under microscopy and amplified. Proteins were extracted and verified with Western Blot to verify for knock-out status. Two colonies with APC knock-out were confirmed. The sgRNA sequences are provided in Table S2.

### Pharmacological inhibitor treatments

Prior to fixation for immunostaining or biochemical analysis, freshly passaged 3D organoids were treated with 10 μM of CHIR for 72 h or 200 nM LGK for 48 h. For live imaging, inhibitors were added 48h (CHIR), 24h (LGK) prior imaging to ensure optimal cell viability. For ECCD-1 treatments, organoids were mixed with 100 μM ECCD-1 prior to embedding in BME2 and cultured with ENR containing 100 μM ECCD-1 for 24h.

### Fluorescence loss in photoactivation (FLIP) assay

Confocal microscopy was performed using a laser scanning inverted fluorescence microscope (Zeiss LSM 510 META, Zeiss, Jena, Germany) equipped with a 40x 1.4 NA plan Apochromat water immersion objective and 37°C temperature control (Zeiss, Jena, Germany). GFP fluorescence was collected with a 505–530 bandpass filter after excitation with a 25-milliwatt argon laser emitting at 488 nm. FLIP experiments were performed by first defining regions of interest (ROIs), which were repeatedly activated using a 583 nm laser while an image was acquired with reduced laser power (0.5% output) at the start of the experiment and after each bleach. Three iterations with 100% laser power were used for the bleaching pulses. An eventual pause between the bleaches ensured some recovery in the ROI. Bleaching time was usually <10 s, resulting in bleaching throughout the full thickness of the junction. After bleaching, images were taken within the same focal plane at regular intervals (between 2 and 30 s) to monitor fluorescence recovery. FLIP curves were fitted using a one phase exponential decay function, *I = (I_0_- I_p_)*exp(−K*t) + I_p_*, where *t* is the time, *I* is the intensity, *I_0_* is the initial intensity value at *t*=0, *I_p_* is the final intensity plateau value and *K* is the decay rate constant). Based on the fits, we determined the mobile fraction as *(I_p_ - I_o_) / (I- I_0_)* with and the decay half-time (t_1/2_) as *ln(2) / K*.

### Particle image velocimetry (PIV) analysis

Live imaging time series were obtained using an AxioObserver SD Spinning Disk microscope as described above for FLIP experiments, with time intervals of 10 min for 24 h. PIV analysis was performed as previously described, based on openpiv^116,117^. For determination of vectors, a window size of 15 μm × 15 μm with 50% overlap was used. Calculation of mean migration velocities and correlation lengths was performed as previously described^40,46^.

### Quantitative real-time PCR (RT-qPCR)

Total RNA samples were prepared from small intestinal organoids using the RNeasy Plus mini kit according to the manufacturer’s instructions. To quantify Wnt targets markers expression, one-step quantitative real-time PCR was performed using the Luna Universal qPCR Master Mix. Quantitative real-time PCR reactions were performed on an Applied Biosystems Quant Studio 7 Flex Real-Time PCR System. The cycling conditions were 1 cycle of denaturation at 60°C for 10 min, then 95°C for 2 min, followed by 39 amplification cycles at 95°C for 10 s and 60°C for 30 s. The delta-delta-cycle threshold (ΔΔCT) was determined relative to GAPDH or TATA-binding protein (Tbp) and control samples. Analysis was performed using Quant Studio Real-Time PCR Software. Primer sequences are provided in Table S2.

### Indirect immunofluorescence staining of 3D organoids and organoid monolayers

Samples were fixed in 4% paraformaldehyde (PFA) in 1x PBS for 30 min (3D organoid culture) or 15 min (2D organoid monolayers) at room temperature and permeabilized using 1% Triton X-100 in PBS for 5 min at room temperature. In the case of immunostaining for NUP98, fixation was performed using 4% PFA + 0.3 % Triton in PBS. Samples were blocked in PBS + 5% BSA + 0.05% Tween-20 (PBS-T) for 1 h at room temperature and incubated with primary antibodies (1:100 in PBS-T, overnight, 4°C; see Table S1) followed by AlexaFluor-conjugated secondary antibodies (1:1000, overnight, 4°C) and DAPI (0.5 µg mL^−1^, 30 min). Phalloidin-AlexaFluor647 (1:2000) was added with secondary antibodies for F-actin counterstaining. Samples were washed thoroughly three times with PBS-T (0.1% Tween-20) between steps. After washing, one drop of Fluoromount-G was added on the top of the sample and the sample was mounted with a coverslip.

### Preparation and indirect immunofluorescence staining of mouse small intestine

Immediately following sacrifice, the small intestine (SI) was removed by incising distal to the pylorus and proximal to the caecum. The mesentery was stripped, luminal contents flushed with ice-cold Ca²⁺/Mg²⁺-free PBS, cut to pieces of 50-60 mm thickness and soaked in 4% PFA in PBS with agitation in the dark for 1 h. After washing 3×15 min with PBS with agitation, the tissue was soaked in 15% sucrose for 1 h, then in 30% sucrose for 2 h. Dehydrated tissue samples were placed in an embedding mold with respect to the appropriate slicing direction and filled quickly with OCT on dry ice and incubated in -20°C for at least 4 h. Embedded tissue was circumferentially sliced using a microtome at 30 μm thickness, deposited on glass slides and preserved directly at -20°C until use. For immunostaining, tissue slices were rehydrated by soaking in PBS containing Ca²⁺/Mg²⁺ for 10 min. Immunostaining was performed using the same protocol as for organoids monolayer above, starting with the permeabilization step.

### Confocal microscopy of immunostained samples

All immunofluorescence samples were imaged using an LSM980 Airyscan 2 microscope (Carl-Zeiss Microscopy, Germany) equipped with a LD LCI Plan-Apochromat 40x/1.2 Water immersion objective. Images were acquired in 4Y multiplex mode using the following diode laser/beamsplitter (detection range) combinations: DAPI channel: 405 nm/300-720nm, green channel: 488 nm/495-550 nm, red channel: 561 nm/573-627 nm, far red channel: 640 nm/574-720 nm. Airyscan raw images were 3D-processed using Zen Blue software (v3.3).

### Western blotting

3D organoids were removed from BME domes as described above for plating organoid monolayers including centrifugation at 500xg for 5 min. Following centrifugation, RIPA buffer supplemented with an EDTA-free protease inhibitor cocktail was added to the pellet to lyse the organoids. The lysate solution was kept on ice for 15 mins before centrifuging at 16000xg for 15 mins at 4°C. The supernatant was removed and protein concentration was determined by Bradford assay. 20 µg protein per lane was resolved by SDS-PAGE, transferred to nitrocellulose, blocked (5% Roti Block in TBS-T), and probed with primary antibodies overnight at 4°C followed by HRP-conjugated secondaries for 2 h at room temperature. Bands were visualized using ECL substrate, imaged using a Fusion Solo documentation station (Vilber) and quantified by densiometry using the Fiji distribution of Image J (henceforth Fiji)^118^. Raw uncut images of the membranes can be found in the supplemental information (Figure S6).

### 3D segmentation of intracellular compartments

Segmentation of intracellular compartments was used to determine relative nuclear, cytoplasmic and junctional intensities for quantification of immunostaining experiments. Samples were immunostained and quantified as described above. To segment cells in 3D, we used an active contours approach, using the segmented nuclei as seeds to identify cellular volumes. Initially, nuclear segmentation in 3D was performed as previously described using a custom model with StarDist^119–121^. Next, we obtained the volume of the monolayer using iterative thresholding and GPU-accelerated binary morphology operations on the Phalloidin counter-stain channel in Fiji with CLIJ2^122^. The binary contour of the monolayer volume was then added to the Phalloidin channel to limit the cellular segmentation to this bound. Segmentation of cellular volumes by active contours was performed using the LimeSeg plugin in Fiji (d_0=”5.0“; f_pressure = “0.025“; iterations = “1000”), using spherical volumes of 10 px centered at the nuclear centers of mass as initial volumes^123^. The resulting segmentations point clouds were then used to generate 16-bit label image and visualized using a custom python script, leveraging the following plugins: vedo^124^, open3d^125^ and napari^126^. Following this step, we generated label images for cell nuclei, membranes and cytoplasm volumes. The membrane volume was generated by taking a volume from 2 px outside of the cell segmentation contour to 1 px inside of the cell segmentation using binary filter operations. A maximum filter was used for label dilation, and a Laplace filter was used for label erosion. The membrane volume was further refined by splitting the volume into four vertical quartiles. The upper quartile was then designated as “apical membrane”, the middle two quartiles as “lateral membrane” and the lower quartile as “basal membrane.”

### 3D segmentation of nuclear cortex regions

To determine the nuclear cortex to nuclear volume intensity ratio for NUP98 immunostaining, 3D nuclear volumes were segmented as described above. The nuclear cortex volume was generated using a custom python script taking a volume from 1 px outside of the nuclear segmentation contour to 2 px inside of the cell segmentation using binary filter operations. The inner nuclear volume was taken as the voxels contained within the cortex.

### Reconstruction of crypt-villus axis in organoid monolayers

To computationally reconstruct the crypt-villus axis in organoid monolayers, binary masks of circular crypt-like compartments were manually generated in Fiji based on visual inspection of the DAPI and Phalloidin counter-stainings. Crypt-like compartments have a distinctly high density of nuclei in the xy plane due to the tall columnar shapes of cells in this compartment. These shapes are easily visible in the 3D Phalloidin stacks, with crypt-like compartments appearing as “hills” in an otherwise flat apical profile. Surrounding the inner crypt region is a 1-2 cell-wide row of transit amplifying cells, which have a distinctly elongated shape parallel to the contour of the compartment.

The 3D cell segmentations were then used to determine the centers of mass of each cell in the monolayer. From these positions, each cell was assigned to the nearest crypt-like region based on Euclidean distance to the crypt centroids. Each cell then received a relative position, x, along the crypt-villus axis to the nearest crypt, and this distance was rescaled with respect to the contour of the crypt such that the distance from the crypt center to the edge of the crypt contour is equivalent to 1. Therefore, all cells within the crypt-like compartment received a rescaled position in the range [0,1], and cells outside of the crypt received a rescaled position × > 1. For quantifications where values are averaged for individual compartments (crypt, TA, villus), the crypt was defined 0 < × < 0.8, the TA zone was defined as 0.8 < × ≤ 1.2 and the villus was defined as 1.2 < ×.

### Quantification of β-catenin and cellular aspect ratio along the crypt-villus axis

To quantify β-catenin intensity along the reconstructed crypt-villus axis, cellular position values were binned with respect to the rescaled positions from 0 to 3, with bin widths = 0.1 in rescaled distance units. For the analysis of β-catenin along the crypt-villus axis in 3D organoids and mouse intestinal slices, cell positions were rescaled in a similar manner, with the base of the crypt or bud region corresponding to position × = 0 and the upper/inner end of the neck of the crypt/bud region corresponding to position × = 1. The intensities of cells within each binned region were then averaged, and the mean ± SEM was plotted in the corresponding linescans.

For measurements of cellular aspect ratio in organoid monolayers, 3D cell segmentations were used. Cellular morphometric properties were analyzed using the MorphoLibJ package in Fiji^127^. The in-plane aspect ratios were defined as the ratio of the two smallest principal radii (R2/R3) from a 3D ellipsoid fit. As these cells are columnar, the principal axis is oriented along the apico-basal axis. Aspect ratio values were binned and averaged along the reconstructed crypt-villus axis as described for the β-catenin intensity measurements.

### Quantification of cellular APC distribution

To quantify the relative apical to basal APC intensity ratio, apical, lateral and basal cell membranes were segmented in 3D as described above. For each cell within the crypt contour, the mean intensity in the apical membrane was divided by the mean intensity in the basal membrane. Total APC intensity was defined as the average intensity in the whole cell segmentation.

### Generation and analysis of TOP-GFP fluorescence linescans

To generate TOP-GFP linescans over time for live imaging data, mask images for each timepoint were generated manually based on the crypt positions given by the arrangement of TOP-GFP^hi^ cells. For each circular crypt region, linescan position coordinates were determined for 360 linescans for each directional degree. As described above, positions were rescaled such that position × = 1 is equivalent to the crypt edge. In this way, linescan positions were determined to extend from × = 0 to × = 1.4, with a total of 100 equidistant points along each linescan. For each linescan point, the intensity was determined by cubic interpolation. Each of the 360 linescans was then aligned and averaged by taking the mean to achieve a single radially averaged linescan at each time point. Any points outside of the crypt region that were not within the image were not used for averaging.

### Lucifer Yellow organoid permeability assay

The organoid-based epithelial permeability assay is based on a previously published protocol^128^. 3D Organoids were passaged as described above, and 25 µL of the organoid BME mixture was dispensed into each compartment of a 4-chamber glass-bottom dish. After allowing the organoids to settle and adhere for 24 hours, the culture medium was replaced with treatment medium as described above. Following treatment, organoids were transferred to the microscope stage and maintained at 37°C with 100% relative humidity and 5% CO2 throughout imaging. Live imaging was performed using a Cell Observer Z1 epifluorescence microscope (Carl Zeiss Microscopy, Germany) equipped with an Axiocam 503 camera and an EC Plan-Neofluar 20x/0.5 objective. Lucifer Yellow fluorescence intensity was acquired using a 470 nm LED light source in combination with a Zeiss filterset 62. Just prior to imaging, Lucifer Yellow was added to the culture medium to a final concentration of 1 mM. Timelapse imaging was performed every 7.5 minutes over a 5-hour period to monitor dye permeability into the organoid lumen. Organoid positions were tracked during the experiment using an automated stage with multi-position imaging. For image analysis, Fiji was used to extract the mean fluorescence intensity of the organoid lumen and background. Relative luminal fluorescence was calculated as the product of the relative lumen fluorescence at each timepoint and its initial value, divided by the product of the background fluorescence and its initial value.

### Additional image analysis

Unless otherwise described above, image analysis was performed using Fiji. Cell height measurements (Figure 4C) were performed using brightfield and Phalloidin-stained immuofluoresence images by manually drawing a line beginning from the actin-brush border to the basal membrane, perpendicular to the contour at the cell-lumen interface. For each organoid ∼10 cells were measured in this manner, and the mean was taken as the cell height value for the organoid. Lumen perimeter roundness (Figure S3B) was assessed using a maximum intensity projection of the phalloidin staining. The lumen contour was drawn manually as indicated in Figure S3A. From the resulting region, the roundness was defined as *(4π A) / P*^2^, where *A* is the region area and *P* is the region perimeter. To determine the relative of TOP-GFP^hi^ to Ki-67+ cell area, a contour surrounding the Ki-67+ cells was drawn manually, and the ratio of the resulting areas was determined. TOP-GFP intensity in 3D organoids (Figure 4E) was assessed by calculating the mean intensity of the entire GFP image over time. The signal was normalized by taking the intensity ratio at t_i_ over t_0_. Luminal volume was estimated by manually drawing the lumen contour from a cross-sectional 2D image slice at each time point. The time scale was shifted to ensure images were aligned such that t_0_ corresponded to the maximum volume, and the estimated volume was normalized by taking the ratio at t_i_ over t_0_. Individual TOP-GFP^hi^ cells were manually tracked in 3D organoids (Figure 4F) and organoid monolayers (Figure 4H) using the MTrack2 plugin in Fiji. Mean instantaneous speed was averaged across all timeframes for each cell being tracked. Persistence was defined as the ratio of the total distance travelled (from the position at the final time point to position at the first time point) over the total path length. To determine the oscillation rate of TOP-GFP^hi^ cells in 3D oragnoids (Figure 4H), oscillations were defined as directional changes with an angle θ > 135 deg. or θ < -135 deg. The oscillation rate was defined as the number of oscillations divided by the total imaging time. To determine the relative area of Ki-67+ and OLFM4+ cells in 3D organoids (Figure 5A), images were manually thresholded to generate a binary mask for the positive cells. The resulting area of this mask was then divided by the area of a manually thresholded mask for the entire organoid. To determine the area of crypts comprising TOP-GFP^hi^ cells (Figure 5C), a contour was manually drawn around the crypts using a brightest point projection of the TOP-GFP image.

### Data visualization and statistical analysis

Generation of boxplots, bar plots and statistical analysis was performed using GraphPad Prism. For box plots, dots represent individual measurements as described in the figure legends. Boxes represent the 25^th^ to 75^th^ percentiles of the data. The line inside of the box is the median. Whiskers extend to the minimum and maximum value. Bonferroni’s post-hoc test is defined as pairwise t-tests using Fisher’s LSD test, which assumes equal variance among samples and is 2-sided. The SD used in this test is the SD pooled across all samples. The Bonferroni correction is applied to determine the reported significance.

For custom image analysis and visualization in python, the miniconda distribution of python was used, and virtual environments for different image and data analysis tasks were generated using conda or pip package managing functions. In addition to the packages listed above for specific tasks, the following packages were used: matplotlib^129^, numpy ^130^, pandas^131^, scikit-image^132^, scipy^133^ and tifffile^134^. The Large Language Model ChatGPT was used to assist in writing custom python functions for image analysis. All code segments generated by ChatGPT were tested and validated.

## Notes

### Competing Interest Statement

The authors have declared no competing interest.

