## Supplemental Information for "Interplay between β-catenin transcriptional and cell-cell junction activity regulates homeostasis and collective dynamics in the intestinal epithelium"

#### Supplemental Figures

##### Figure S1

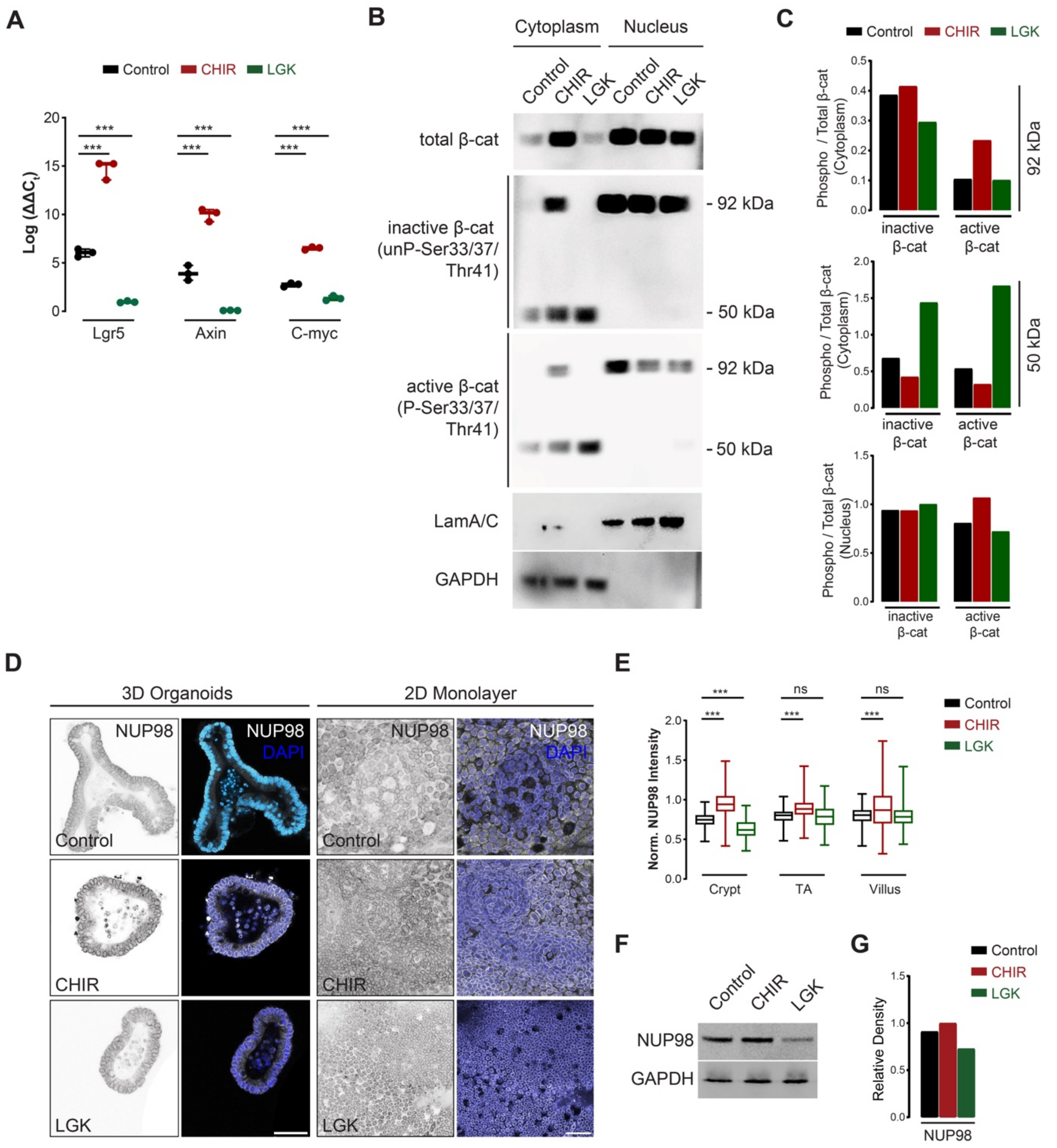


**Figure S1. Modulation of Wnt signaling influences downstream signaling and β-catenin nuclear import. A.** RT-qPCR of Wnt pathway targets following CHIR/LGK treatment in 3D organoids. **B.** Western blot analysis of expression, post-translational modification and sub-cellular localization of β-catenin following cellular fractionation to obtain cytoplasmic (GAPDH+) and nuclear (LaminA/C+) fractions for organoids treated with CHIR or LGK. **C.** Quantification of Western blot in *B* by densiometric analysis. Intensity levels were normalized to the GAPDH loading control (cytoplasmic) or LaminA/C (nuclear). **D.** Representative micrographs of organoids in 3D culture or monolayer culture treated with CHIR or LGK and immunostained for NUP98 and counterstained with DAPI. SB: 50μm, 50μm. **E.** Quantification of relative intensity of NUP98 at the nuclear cortex (with respect to the inner nucleus volume) in organoid monolayers as for conditions as in *D*. Data is compiled $n = 12, 15$ monolayers and $N = 3, 3$ independent experiments. **F.** Western blot of β-catenin and NUP98 for organoids treated with CHIR or LGK. GAPDH was used as a loading control. **G.** Quantification of Western blot in *F* by densiometric analysis. Intensity levels were normalized to the GAPDH loading control. Statistical tests performed using one-way ANOVA tests ($p<0.001$ for all plots) and Bonferroni's post-hoc test (***$p<0.001$, ns:$p>0.05$).

**Figure S2**

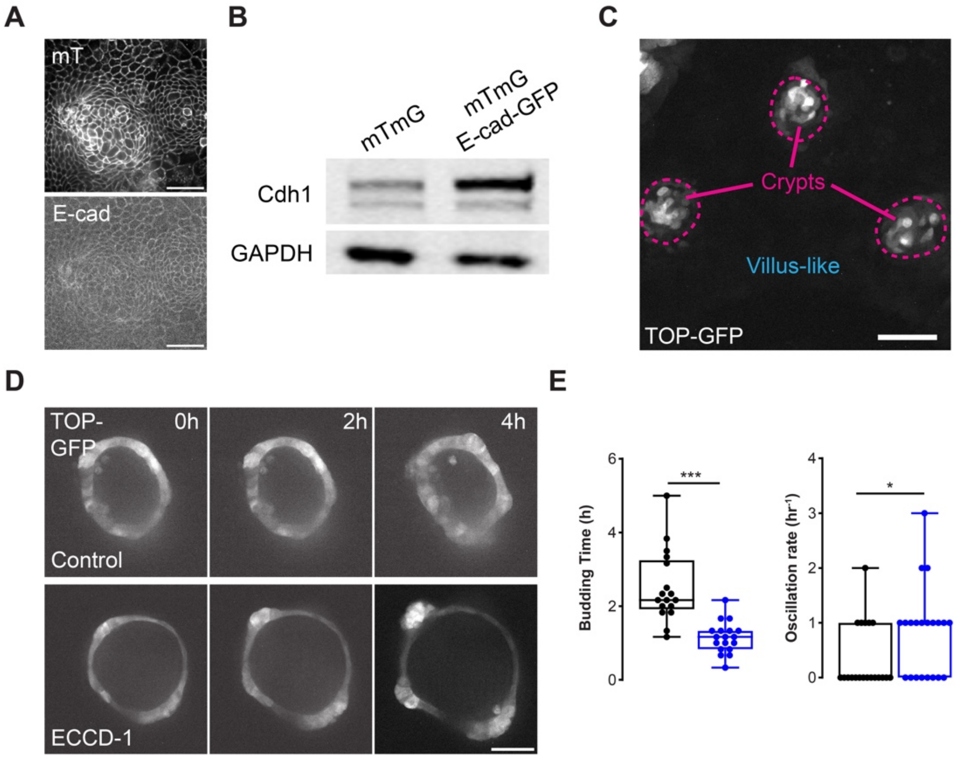


**Figure S2. Validation of E-Cadherin overexpressing organoids and effects of E-cadherin blocking on organoid growth dynamics. A.** Representative fluorescence micrographs of mTmG/E-cad-GFP (Cdh1-GFP) organoid monolayer. Cdh1-GFP is expressed from a bacterial chromosome (BAC) to induce a slight overexpression. SB: 50 μm. **B.** Western blot analysis of Cdh1 expression in parental mTmG and mTmG/E-cad-GFP organoids. GAPDH is used as a loading control. **C.** Representative micrograph of TOP-GFP organoid monolayer, indicating several crypt-like compartments enriched in TOP-GFP$^{hi}$ stem cells (magenta dotted lines). SB: 50 μm. **D.** Representative micrographs of 3D TOP-GFP organoids treated with ECCD-1 for 24 hours prior to beginning live imaging. SB: 50 μm. **E.** Quantification of time to first bud formation and oscillation rate of TOP-GFP$^{hi}$ cells during bud specification as for conditions in *D* from $n = 24$, 24 organoids and $N = 3$, 3 independent experiments. Statistical tests performed using Welch's t-test (*p<0.05, ***p<0.001).

**Figure S3**

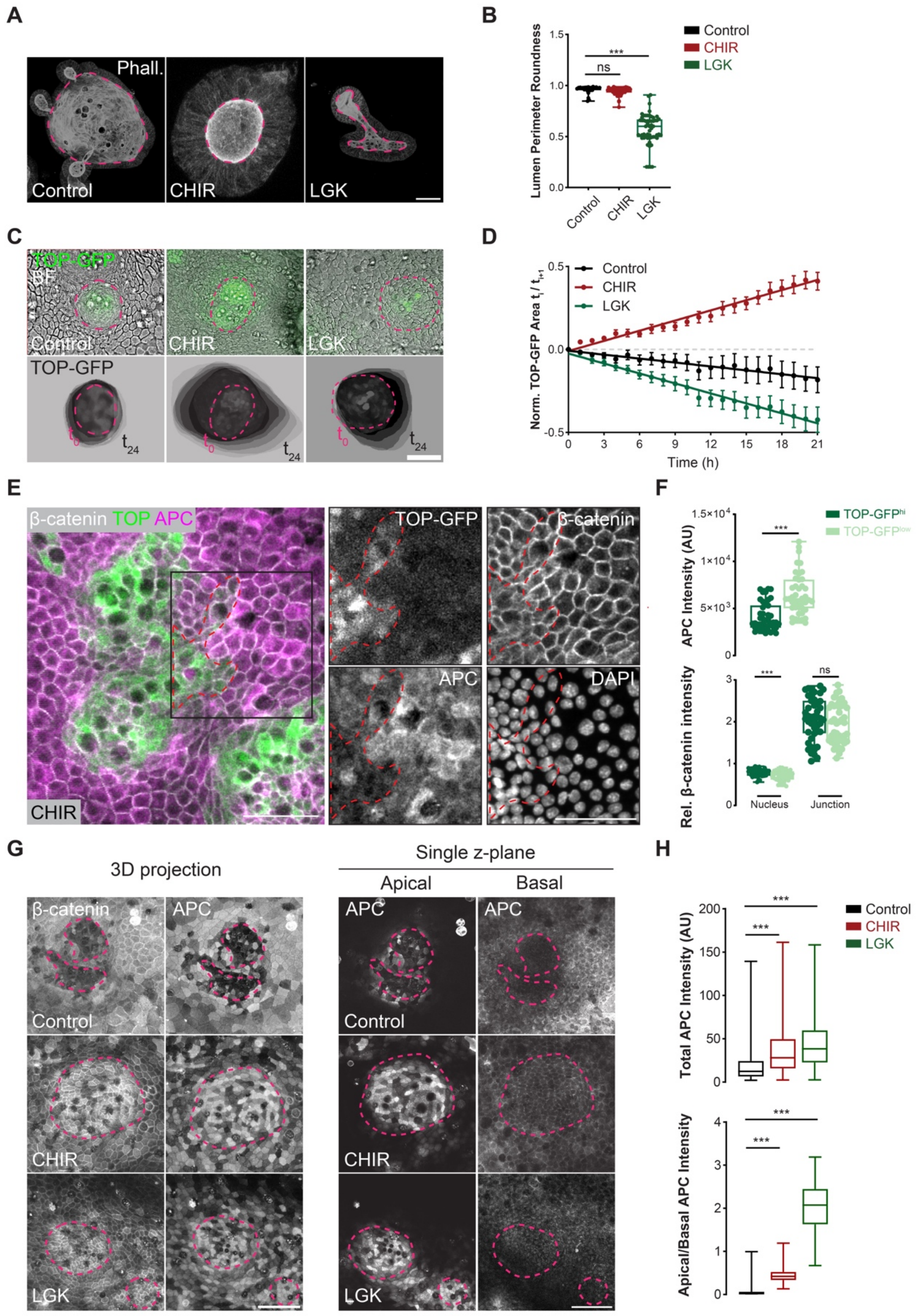


**Figure S3: Modulation of Wnt signaling disrupts tissue compartmentalization and localization of destruction complex components. A.** Representative micrographs of 3D organoids treated with CHIR or LGK and stained with fluorescent Phalloidin (Phall.) to label

F-Actin. SB: 50 µm. **B.** Quantification of the roundness of the inner lumen for conditions in *A*. Each dot represents one 3D organoid from *n* = 40, 40, 40 organoids and *N* = 4, 4, 4 independent experiments. **C.** Representative micrographs of TOP-GFP organoid monolayers treated with CHIR or LGK. *Lower panel*, overlay of crypt morphology over time from a cropped region of the upper panel around the indicated crypt-like compartment. The scale is equivalent in the top and bottom panels; region of the initial time point, $t_0$, is shifted to be aligned with the center of the frame. SB: 50 µm. **D.** Quantification of area of the crypt-like compartment over time as for conditions in *C*. Data is normalized and shifted such that the area of the first time point is 0. Dots represent mean ± SEM. Lines represent linear fits of area over time for each condition. Data compiled from *n* = 12, 12, 12 organoid monolayers and *N* = 3, 3, 3 independent experiments. **E.** Representative micrographs of TOP-GFP organoid monolayers treated with CHIR or LGK and immunostained for β-catenin and APC and counterstained using DAPI. Images are maximum projections of z-stacks. Image panels on the right correspond to the region indicated by the black box in the merge image. Red dashed lines indicate TOP-GFP$^{hi}$ cells migrating out of the crypt-like region. SB: 50 µm, 50 µm. **F.** Quantification of APC and β-catenin intensity within the TOP-GFP$^{hi}$ region indicated by the red dashed line and the surrounding TOP-GFP$^{low}$ cells (as indicated in the example in *E*). β-catenin localization is segregated into relative nuclear and junctional intensity. Each dot represents one cell from *n* = 33 crypts from *N* = 3 independent experiments. **G.** Representative micrographs of organoid monolayers treated with CHIR or LGK and immunostained for β-catenin and APC. Images are displayed as maximum projections of z-stacks (left panel) and single apical and basal planes (right panel). SB: 50 µm, 50 µm. Magenta dashed lines indicate the crypt-like compartments. **H.** Quantification of total and relative apical-to-basal intensity ratio of APC for conditions in *G*. Data compiled from n = 9, 9, 9 and N = 3, 3, 3 independent experiments. Statistical tests performed using one-way ANOVA tests ($p<0.001$ for all plots) and Bonferroni's post-hoc test (***$p<0.001$, ns:$p>0.05$).

#### Figure S4

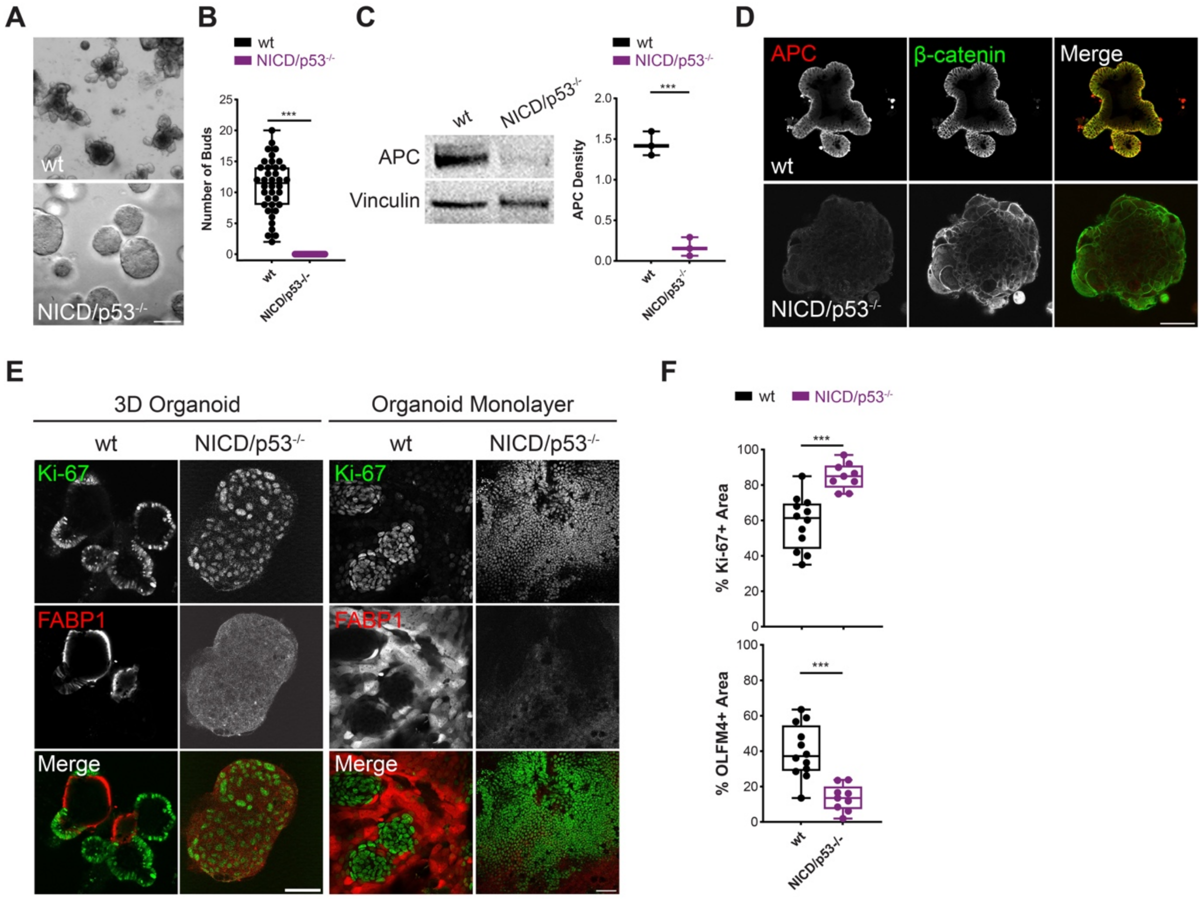


**Figure S4. Mouse small intestinal tumor organoids display perturbed differentiation and crypt/villus organization. A.** Representative brightfield micrographs of wild type (wt) and tumor (NICD/p53$^{-/-}$) organoids in 3D culture, 7 days after passage. SB: 50 µm. **B.** Quantification of number of buds from conditions in *A* from n = 45, 50 organoids from N = 3, 3 independent experiments. **C.** Western blot analysis of APC expression in wt and tumor organoids. Vinculin is used as a loading control. Quantification reflects relative APC signal relative to loading control and corrected for background. Each dot represents one of N=3 independent Western blot experiments. **D.** Representative micrographs of wt and NICD/p53$^{-/-}$ tumor organoids in 3D culture or monolayer culture immunostained for β-catenin and APC. SB: 50µm. **E.** Representative micrographs of healthy and tumor organoids in 3D culture or monolayer culture immunostained for Ki-67 and FABP1. SB: 50 µm, 50 µm. **F.** Quantification of Ki-67 and FABP1 positive area relative to total area for organoid monolayers as in *E*. Each dot represents one organoid monolayer from *n* = 15, 15 monolayers and *N* = 3, 3 independent experiments. Statistical tests performed using Welch's t-test (***p<0.001).

**Figure S5**

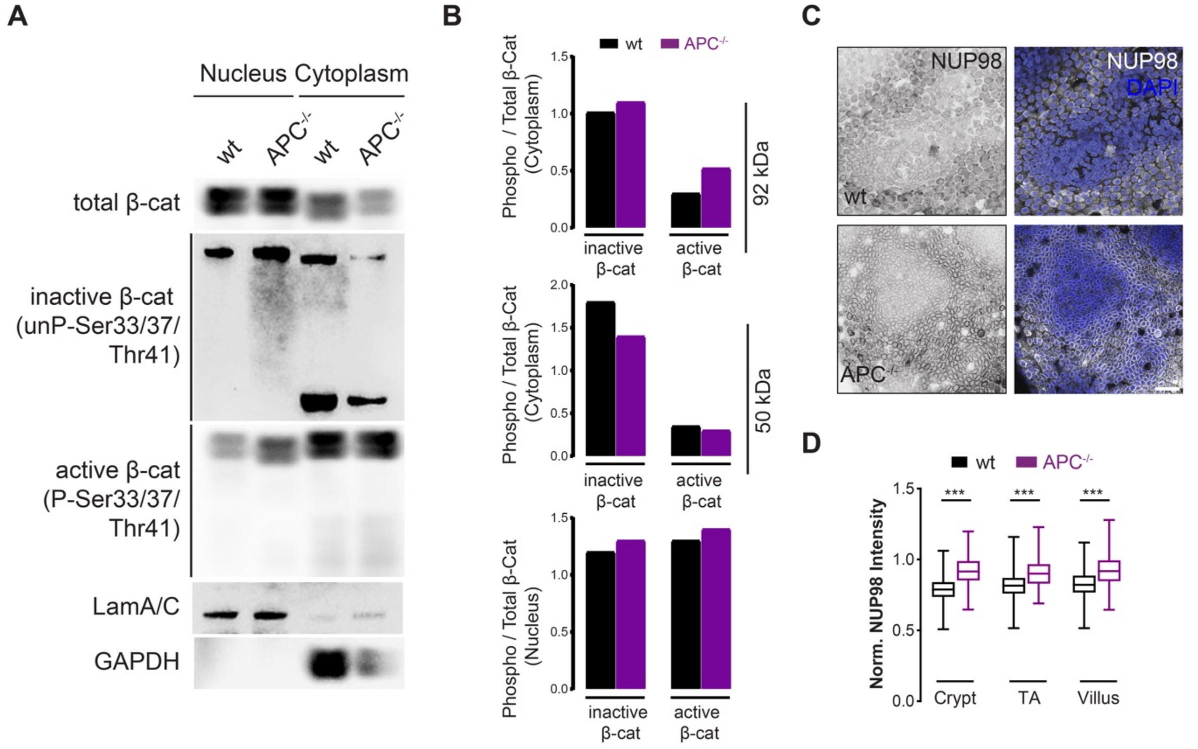


**Figure S5. Analysis of sub-cellular localization of β-catenin in wild type and APC$^{-/-}$ organoids.** **A.** Western blot analysis of expression and sub-cellular localization of β-catenin and NUP98 in healthy wild type (wt) and APC$^{-/-}$ organoids following cellular fractionation to obtain cytoplasmic (GAPDH+) and nuclear (LaminA/C+) fractions. **B.** Quantification of Western blot in *A* by densiometric analysis. Intensity levels were normalized to the GAPDH or LaminA/C loading controls. **C.** Representative micrographs of wt or APC$^{-/-}$ organoid monolayers immunostained for NUP98 and counterstained with DAPI. SB: 50µm. **D.** Quantification of relative intensity of NUP98 at the nuclear cortex (with respect to the inner nucleus volume) in organoid monolayers as for conditions as in *C*. Data is compiled from *n* = 15, 15 monolayers and *N* = 3, 3 independent experiments. Statistical tests performed using one-way ANOVA tests (p<0.001) and Bonferroni's post-hoc test (***p<0.001).

**Figure S6**

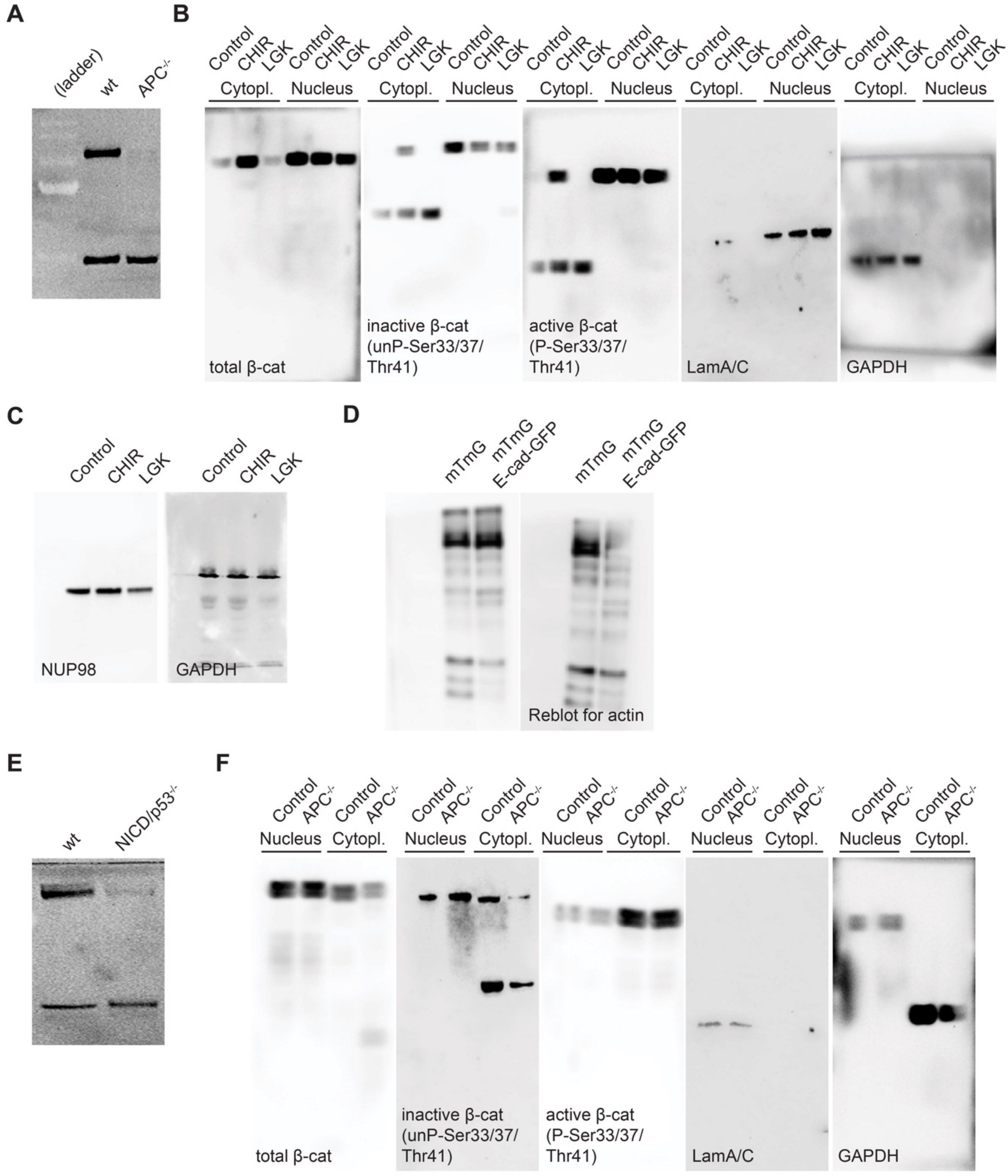


**Figure S6. Uncropped Western blots. A.** PDVF membrane following Western Blot showing protein bands corresponding APC and Vinculin in wild type (wt) and APC$^{-/-}$ organoids (Fig. 6B). **B.** Nitrocellulose membrane following Western Blot showing protein bands corresponding total β-catenin, inactive β-catenin, active β-catenin, Lamin A/C and GAPDH as controls for fractionation of nucleus and cytoplasm, respectively (Fig. S1B). **C.** Nitrocellulose membrane following Western Blot showing protein bands corresponding NUP98 and actin (Fig. S1F). **D**.

Nitrocellulose membrane following Western Blot showing protein bands corresponding to E-cadherin (Cdh1) and actin (Fig. S2B). **E.** PDVF membrane following Western Blot showing protein bands corresponding to APC and Vinculin (Fig. S4F). **F.** Nitrocellulose membrane after Western Blot showing protein bands corresponding β-catenin total, inactive β-catenin, active β-catenin, LaminA/C and GAPDH as controls for fractionation of nucleus and cytoplasm, respectively (Fig. S5A).

### Supplemental Tables

**Table S1. Critical reagents used in experimental procedures.**

| Reagent | Source | Product Number |
|---|---|---|
| **Antibodies/Fluorescent Dyes** | | |
| Rabbit-α-Olfm4 | Cell Signaling Technologies | 39141 |
| Mouse ECCD-1 | Takarabio | M107 |
| Mouse-α-Ecadherin | Thermo Fisher Scientific | A15784 |
| Rabbit-α-beta-catenin | Thermo Fisher Scientific | 13-8400 |
| Rabbit-α-Phospho-β-Catenin (Thr41/Ser45) | Cell Signaling Technologies | 9565 |
| Rabbit-α-Non-phospho (Active) β-Catenin (Ser45) | Cell Signaling Technologies | 19807 |
| Rabbit -α-APC | Thermo Fisher Scientific | PA5-119648 |
| GFP-Booster Alexa 488 | ChromoTek | gb2AF488 |
| Rabbit-α-NUP98 (C39A3) | Cell Signaling Technologies | 2598 |
| Mouse -α-Ki67 | BD Sciences | 393778 |
| Rabbit -α-FABP1 | Cell Signaling Technologies | 13368 |
| Goat-α-Mouse-IgG-AlexaFluor488 | Thermo Fisher Scientific | A-11001 |
| Goat-α-Rabbit-IgG-AlexaFluor568 | Thermo Fisher Scientific | A-11011 |
| Goat-α-Mouse-IgG-AlexaFluor568 | Thermo Fisher Scientific | A-11004 |
| Goat-α-Rabbit-IgG-AlexaFluor647 | Thermo Fisher Scientific | A-21244 |
| Goat-α-Mouse-IgG-AlexaFluor647 | Thermo Fisher Scientific | A-21235 |
| Goat-α-Rabbit-IgG-HRP | Merck, Sigma-Aldrich/Chemicon | AP307P |
| Goat-α-Mouse-IgG-HRP | Merck, Sigma-Aldrich/Chemicon | AP308P |
| 4,6-diamidino-2-phenylindole (DAPI) | Thermo Fisher Scientific | D1306 |
| Phalloidin AlexaFluor647 | Thermo Fisher Scientific | A22287 |
| Lucifer Yellow | Thermo Fisher Scientific | L453 |
| **Chemicals, peptides, and recombinant proteins** | | |
| $PBS^{-/-}$ (1x DPBS without $Ca^{+2}/Mg^{+2}$) | Thermo Fisher Scientific, Gibco | 14190 |
| $PBS^{+/+}$ (1x DPBS with $Ca^{+2}/Mg^{+2}$) | Thermo Fisher Scientific, Gibco | 14040 |
| DMEM/F-12 | Thermo Fisher Scientific, Gibco | 10565018 |
| Cultrex BME | Bio-Techne, R&D systems | 3533-010-02 |
| Culturex Organoid Harvesting Solution | Bio-Techne, R&D systems | 3700-100-01 |
| Fetal Calf Serum (FCS) | PAN-Biotec | P30-3031 |
| Gibco Antibiotic/Antimycotic (100X) | Thermo Fisher Scientific, Gibco | 15240096 |
| GlutaMAX-1 (100X) | Thermo Fisher Scientific, Gibco | 35050061 |
| Murine EGF | Thermo Fisher Scientific, PeproTech | 315-09 |
| Murine FGF | Thermo Fisher Scientific, PeproTech | 450-33A |
| N2 Supplement (100X) | Thermo Fisher Scientific, Gibco | 17502048 |

| | | |
|---|---|---|
| B27 Supplement (50X) | Thermo Fisher Scientific, Gibco | 17504044 |
| Y-27632 dihydrochloride | TebuBio | 282T1725 |
| Ciprofloxacin | Merck, Sigma-Aldrich | 17850-5G-F |
| Metronidazol | Merck, Sigma-Aldrich | M1547 |
| DMSO | Carl Roth | 4720.4 |
| Trypsin/EDTA 0,5% 10x | Thermo Fisher Scientific, Gibco | 15400054 |
| 40% Acrylamide Solution | Bio-Rad | 1610144 |
| 2% Bis-Solution | Bio-Rad | 1610142 |
| Ammonium Persulfate | Merck, Sigma-Aldrich | A7460 |
| N,N,N',N'-tetramethylethylenediamine | Merck, Sigma-Aldrich | 1107320100 |
| FluoSpheres (0.2 µm, yellow-green) | Thermo Fisher Scientific, Invitrogen | 10513463 |
| 16% Formaldehyde (w/v) | Thermo Fisher Scientific | 28908 |
| Glutaraldehyde | Merck, Sigma-Aldrich | G6257 |
| (3-Aminopropyl) trimethoxysilane | Merck, Sigma-Aldrich | 440140 |
| Laminin 1 (Mouse) | Thermo Fisher Scientific | 23017015 |
| Rat Tail Collagen Type 1 | Corning | 354236 |
| HEPES (1M) | Thermo Fisher Scientific, Gibco | H3537 |
| Pluronic F-127 | Merck, Sigma-Aldrich | P2443 |
| Sulfo-SANPAH (Sulfosuccinimidyl-6-4-azido-2'- nitrophenylamino) hexanoat) | Merck, Sigma-Aldrich | 803332 |
| RNeasy Plus mini kit | Qiagen | 74134 |
| Luna Universal qPCR Master Mix | New England Biolabs | M3003 |
| Triton X-100 | Merck, Sigma-Aldrich | T8787 |
| Tween-20 | Carl Roth | 9127.8 |
| Complete protease inhibitor cocktail | Merck, Roche | 11697498001 |
| PhosphoSTOP phosphatase inhibitor | Merck, Roche | 4906845001 |
| ECL Western Blotting Substrate | Thermo Fisher Scientific, Pierce | 32109 |
| Alt-R™ S.p. dCas9 Protein V3 | IDT | 1081066 |
| Transdux | System Biosciences | LV850A-1 |

**Table S2. Oligomer Sequences for sgRNAs and RT-qPCR**

| | | |
|---|---|---|
| sgAPC-RNA (IDT) | Mm.Cas9.APC.1.AA | 5'- mU*mC*mG* rUrGrA rUrCrC rArCrA rCrGrU rGrUrA rGrCrG rUrUrU rUrArG rArGrC rUrArG rArArA rUrArG rCrArA rGrUrU rArArA rArUrA rArGrG rCrUrA rGrUrC rCrGrU rUrArU rCrArA rCrUrU rGrArA -3' |
| | Mm.Cas9.APC.1.AB | 5'- mU*mU*mU* rGrArG rCrGrU rArGrU rUrUrC rArCrU rCrCrG rUrUrU rUrArG rArGrC rUrArG rArArA rUrArG rCrArA rGrUrU rArArA rArUrA rArGrG rCrUrA rGrUrC rCrGrU rUrArU rCrArA rCrUrU rGrArA rArArA -3' |
| | Mm.Cas9.APC.1.AC | 5'- mA*mG*mA* rCrArU rGrArC rArArG rArCrG rGrCrA rGrCrG rUrUrU rUrArG rArGrC rUrArG rArArA rUrArG rCrArA rGrUrU rArArA rArUrA rArGrG rCrUrA rGrUrC rCrGrU rUrArU rCrArA rCrUrU rGrArA rArArA rGrUrG rGrCrA rCrCrG rArGrU rCrGrG rUrGrC mU*mU*mU* rU -3' |
| Primers (Biomers) | Axin 2 | Fw: 5'-ATG GAG TCC CTC CTT ACC GCA T -3'<br>Rev: 5'-GTT CCA CAG GCG TCA TCT CCT T -3' |
| | Lgr5 | Fw: 5'-AGA GCC TGA TAC CAT CTG CAA AC -3'<br>Rev: 5'-TGA AGG TCG TCC ACA CTG TTG C -3' |
| | C-myc | Fw: 5'- CGC TCC AGT TTC CTG AAC CTC A-3'<br>Rev: 5'- ACC AGG AGC TTG GAC TCA TCA G -3' |

#### Supplemental Video Legends

**Video S1.** Live confocal microscopy (single z-slice) of mouse intestinal organoid monolayers expressing β-catenin-mEos2 without treatment (Control) or treated with CHIR or LGK and imaged in the photoactivatable (583 nm) channel. Between timepoints 2 and 3, a 5 μm x 2 μm ROI is activated centered at a point along a cell-cell junction (related to Fig. 1E). Time in MM:SS. SB: 5 μm.

**Video S2.** Live confocal microscopy (single z-slice) of membrane targeted mTomato-expressing (mTmG) organoid monolayers with particle image velocimetry (PIV) analysis, without treatment (Control) or treated with CHIR or LGK (related to Fig. 2A). Time in HH:MM. Scale Vector: 12 μm/h.

**Video S3.** Live confocal microscopy (single z-slice) of mouse intestinal organoids in 3D culture without treatment (Control) or treated with CHIR or LGK. Lucifer Yellow is included in the culture medium to measure epithelial barrier permeability (related to Fig. 2E). Time in HH:MM. SB: 50 μm.

**Video S4.** Live brightfield microscopy of organoid monolayers with particle image velocimetry (PIV) analysis, without treatment (Control) or treated with ECCD-1 (related to Fig. 3D). Time in HH:MM. Scale Vector: 12 μm/h.

**Video S5.** Live confocal microscopy (single z-slice) of TOP-GFP mouse intestinal organoid monolayers without treatment (Control) or treated with ECCD-1. Colored markers and tracks indicate individual cell trajectories (related to Fig. 3F). Time in HH:MM. SB: 50 μm.

**Video S6.** Live confocal microscopy (single z-slice) of TOP-GFP mouse intestinal organoids in 3D culture without treatment (Control) or treated with ECCD-1 (related to Fig. S2D). Time in HH:MM. SB: 50 μm.

**Video S7.** Live confocal microscopy (single z-slice) of TOP-GFP mouse intestinal organoids in 3D culture without treatment (Control) or treated with CHIR or LGK (related to Fig. 4D). Time in HH:MM. SB: 50 μm.

**Video S8.** Live confocal microscopy TOP-GFP mouse intestinal organoids in 3D culture without treatment (Control) or treated with CHIR or LGK. The video reflects the same example as shown in Video S1, but with the image unwrapped to linearize the 3D equatorial contour of the organoid (related to Fig. 4F). Time in HH:MM. SB: 50 μm.

**Video S9.** Live confocal microscopy (single z-slice) of TOP-GFP mouse intestinal organoid monolayers without treatment (Control) or treated with CHIR or LGK (related to Fig. 4H). Time in HH:MM. SB: 50 μm.

**Video S10.** Live brightfield microscopy of wild type (wt) or $APC^{-/-}$ organoid monolayers with particle image velocimetry (PIV) analysis (related to Fig. 6H). Time in HH:MM. Scale Vector: 12 μm/h.

**Video S11.** Live confocal microscopy (single z-slice) of wild type (wt) or $APC^{-/-}$ organoids in 3D culture. Lucifer Yellow is included in the culture medium to measure epithelial barrier permeability (related to Fig. 6J). Time in HH:MM. SB: 50 μm.
